# Interactions between anthropogenic stressors and recurring perturbations mediate ecosystem resilience or collapse

**DOI:** 10.1101/2022.03.27.485937

**Authors:** DA Keith, DH Benson, IRC Baird, L Watts, CC Simpson, M Krogh, S Gorissen, JR Ferrer-Paris, TJ Mason

## Abstract

Insights into declines in ecosystem resilience, their causes and effects, can inform pre-emptive action to avoid ecosystem collapse and loss of biodiversity, ecosystem services and human well-being. Empirical studies of ecosystem collapse are rare and hampered by ecosystem complexity, non-linear and lagged responses, and interactions across scales. We investigated how an anthropogenic stressor could diminish ecosystem resilience to a recurring perturbation by altering a critical ecosystem driver. We studied groundwater-dependent, peat-accumulating, fire-prone wetlands in southeastern Australia. We hypothesised that underground mining (stressor) reduced resilience of these wetlands to landscape fires (perturbation) by diminishing groundwater, a key ecosystem driver. We monitored soil moisture as an indicator of ecosystem resilience during and after underground mining and, after a landscape fire, we compared the responses of multiple state variables representing ecosystem structure, composition and function in wetlands within the mining footprint to unmined reference wetlands. Soil moisture showed very strong evidence of decline without recovery in mined swamps, but was maintained in reference swamps through eight years. Relative to burnt reference swamps, burnt and mined swamps showed greater loss of peat via substrate combustion, reduced cover, height and biomass of regenerating vegetation, reduced post-fire plant species richness and abundance, altered plant species composition, increased mortality rates of woody plants, reduced post-fire seedling recruitment, and local extinction of a hydrophilc fauna species. Mined swamps therefore showed strong symptoms of post-fire ecosystem collapse, while reference swamps regenerated vigorously. We conclude that an anthropogenic stressor may diminish the resilience of an ecosystem to recurring perturbations, predisposing it to collapse. Avoidance of ecosystem collapse hinges on early diagnosis of mechanisms and preventative risk reduction. It may be possible to delay or ameliorate symptoms of collapse or to restore resilience, but the latter appears unlikely in our study system due to fundamental alteration of a critical ecosystem driver.

## Introduction

Ecosystem collapse signals a qualitative change in ecosystem properties including substantial and lasting loss or displacement of biota and re-organization of structure and ecological processes ((Keith, et al., 2013); (Cumming & Peterson, 2017); (Bland, et al., 2018)). The mechanisms that drive such transformations are diverse, varying from deterministic forcing to positive feedbacks ((Dakos, Carpenter, van Nes, & Scheffer, 2015) (Cumming & Peterson, 2017)), as are the temporal patterns of change which vary from smooth to abrupt (Bergstrom, et al., 2021). Examples include the drying of the Aral Sea and its replacement by hypersaline lakes and ephemeral grasslands (Micklin & Aladin, 2008), regime shifts between clear and turbid states in shallow lakes (Carpenter, 2003), displacement of tropical forests by pasture and plantation systems (Hansen, et al., 2013), desertification of grassy rangelands (Bestelmeyer, Duniway, James, Burkett, & Havsted, 2013) and collapse of numerous pelagic marine fisheries and benthic ecosystems (de Young, et al., 2008). Regime shifts are a specific group of mechanisms among the diverse expressions of ecosystem collapse (Keith, et al., 2013), their hallmarks including incremental environmental change, positive feedback mechanisms, hysteresis and sudden transitions into alternative states that are difficult to reverse (Scheffer, Carpenter, Foley, Folke, & Walker, 2001).

The qualitative structural, compositional and functional changes associated with ecosystem collapse have important implications for conserving biodiversity and maintaining ecosystem services, the dual global imperatives mandated under the United Nations Convention on Biological Diversity and Sustainable Development Goals ((United Nations, 1992); (Nations, 2015)). The consequences of ecosystem collapse for human well-being extend across all sectors from health to economic prosperity (Díaz, et al., 2019).

An understanding of the pathways and mechanisms of ecosystem collapse and how they might be mitigated is imperative to successful conservation strategies and actions, yet these mechanisms are generally complex and poorly understood ((Peterson, Carpenter, & Brock, 2003); (Keith, et al., 2015)). (Cumming & Peterson, 2017) note that “Mechanistic theories of collapse that unite structure and process can make fundamental contributions to solving global environmental problems.” Recent decades have seen significant advances in development and testing of early warning indicators to signal when ecosystems may be approaching collapse (Scheffer, et al., 2009). Much of this work aims to detect declines in ecosystem resilience as a causal agent and precursor to collapse (Fig. 1).

**Figure 1.**
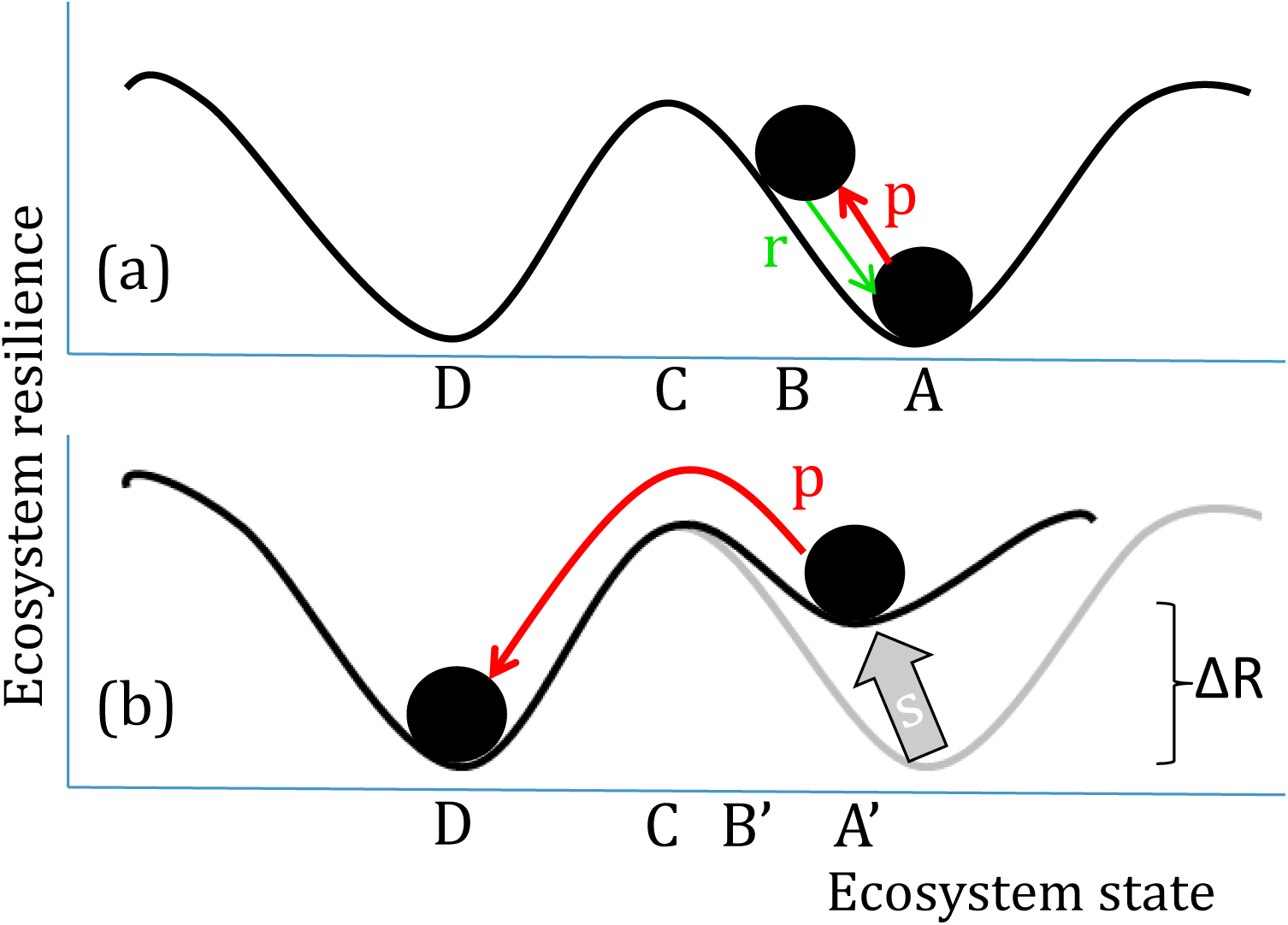
Postulated responses of ecosystem resilience and ecosystem collapse under scenarios of stress and perturbation. In scenario (a), perturbation p results in a transient shift from state A to B, without reaching tipping point, C. The system returns to state A through autogenic recovery, r. In scenario (b), a stressor s diminishes resilience of the system by ΔR and in doing so, shifts the system from state A to A’. When perturbation p occurs, it causes a transformational shift from state A’ beyond tipping point C to collapsed state D.

Methods for early warning of ecosystem collapse have developed from theory and simple models that produce dense time series (Scheffer, et al., 2009). Potential early warning signals include: 1) critical slowing down (CSD) expressed as increasingly slow recovery from perturbations, increased temporal autocorrelations, and increased variance in fluctuations; 2) increasing asymmetry of fluctuations expressed as a skewed distribution of states; 3) ‘flickering’ if stochastic forcing moves the system back and forth between two stable states after it enters the bistable region between them; and 4) spatial signals such as increased spatial coherence among spatial units and other specific patterns that depend on the scale of local drivers (Scheffer, et al., 2009).

If diagnosed and detected, declines in resilience provide an early warning to design and implement adaptive strategies for risk reduction. In this study, we investigated how an anthropogenic stressor, acting through a critical ecosystem driver, could diminish the resilience of an ecosystem to a recurring perturbation, predisposing the ecosystem to an elevated risk of collapse (Fig. 1). Conversely, we hypothesise that the ecosystem is more likely to recover from perturbations if its resilience is not first diminished by the stressor.

Case studies and pathologies have been instrumental to developing theory on ecosystem resilience and collapse ((Holling, 1973); (Scheffer, Carpenter, Foley, Folke, & Walker, 2001); (Cumming & Peterson, 2017); (Bergstrom, et al., 2021)), as have mathematical models derived from systems resilience theory ((Scheffer, et al., 2009); (Dakos, Carpenter, van Nes, & Scheffer, 2015)). Yet, robust empirical studies examining the mechanisms of ecosystem resilience and collapse are extremely rare ((Scheffer, Carpenter, Foley, Folke, & Walker, 2001)). The difficulties of investigation relate to ecosystem complexity, unpredictability and non-linearity of change, ecological lags in responses, large organizational scales and interactions across scales, as well as logistic challenges of replication and definition of suitable experimental controls or reference systems.

Our study system is a type of groundwater-dependent, peat-accumulating wetland ecosystem (known locally as upland swamps) in a landscape with a long history of recurring landscape fires (a perturbation regime). We hypothesised that hydrological change caused by underground mining (an anthropogenic stressor) diminishes the resilience of these wetlands to fires. We first described the postulated mechanism of collapse, and then examined empirical evidence that underground mining initiates changes in groundwater hydrology, a key driver of the wetland ecosystem. We then measured a wide range of state variables as the ecosystem regenerated after a wildland fire to examine differences in response between wetlands exposed to underground mining relative to reference systems located beyond the mining footprint. Finding strong evidence in support of our model, we suggest that this mechanism of ecosystem collapse may be more pervasive than previously appreciated. We review the consequences and irreversibility of collapse in these ecosystems and identify management strategies for impact avoidance. Finally, we consider the broader ecosystem management implications for early detection and risk reduction strategies, given ecological lags between the initiation of the stressor and the transition to collapse.

## Methods

### Study system

We studied ecosystem resilience and collapse in geographically restricted peat-accumulating, groundwater-dependent palustrine wetlands (hereafter upland swamps) in the upper Blue Mountains within the Sydney stratigraphic basin, centred on lat. 33° 23’ S, longitude 150° 13’ E, southeastern Australia. These peaty wetlands are listed as Endangered Ecological Communities under national legislation as ‘Temperate Highland Peat Swamps on Sandstone’ (DEWHA, 2005) and in NSW as ‘Newnes Plateau Shrub Swamps’ (NSW Scientific Committee, 2005). They are treeless ecosystems within a eucalypt forest matrix and characterised by dense growth of hydrophilic sclerophyll shrubs and graminoids, with many hydrophilic specialists and some local endemics represented in the flora and fauna (Benson & Baird, 2012). High levels of subsoil moisture are sustained by perched aquifers on friable sandstones of the Triassic Narrabeen Group, interbedded with low-permeability claystone members (Benson & Baird, 2012). Basal dates of sediments suggest the peatlands developed during the Pleistocene-Holocene transition when the climate became warmer and wetter (Chalson & Martin, 2009); (Young, 2017)). Charcoal throughout the peaty sediments indicates a long history of fire (Chalson & Martin, 2009).

Sediments and vegetation function as landscape sponges that retain, filter and slowly release high quality water, even during prolonged dry periods (Cowley, Fryirs, & Hose, 2018). These landscape hydrological functions are linchpins of ecosystem services to Sydney’s largest city and surrounding regions, including provision of potable water, recreational resources, carbon sequestration and flood mitigation (e.g. (Cowley & Fryirs, 2020); Cowley et al. 2020). Coal is extracted from the Lithgow Coal Seam 200-400 m below the surface of the Newnes plateau, with mining leases currently covering about two-thirds of the swamp distribution.

### Model of ecosystem resilience and collapse

Three environmental conditions are critical to formation and persistence of upland swamps ((Keith, Elith, & Simpson, Predicting distribution changes of a mire ecosystem under future climates, 2014); (Young, 2017)): i) a humid climate (with precipitation exceeding evapotranspiration); ii) low topographic relief (hence slow runoff rates); and iii) low substrate permeability (inhibiting percolation and promoting perched aquifer development).

Fires consume standing vegetation, liberate resources, stimulate regenerative processes and may locally consume peaty substrates depending on their pre-fire moisture content ((Prior, French, Storey, Williamson, & Bowman, 2020)). Functional upland swamps have a resilient response to fire as (Fig. 2), returning to their pre-fire ‘basin of attraction’ through autogenic processes (Folke, et al., 2004). Resilience of the system rests on positive feedbacks in which abundant subsoil moisture limits combustion of peat, promotes dense vegetation that traps sediment, interrupting water flow and maintaining a moist, shady microclimate, limiting entry of non-hydrophiles into the system (Young, 2017). Rapid post-fire regrowth enables upland swamps to function as refuges for fauna and hydrophilic flora in the burnt landscape amidst the more slowly regenerating forest matrix.

**Figure 2.**
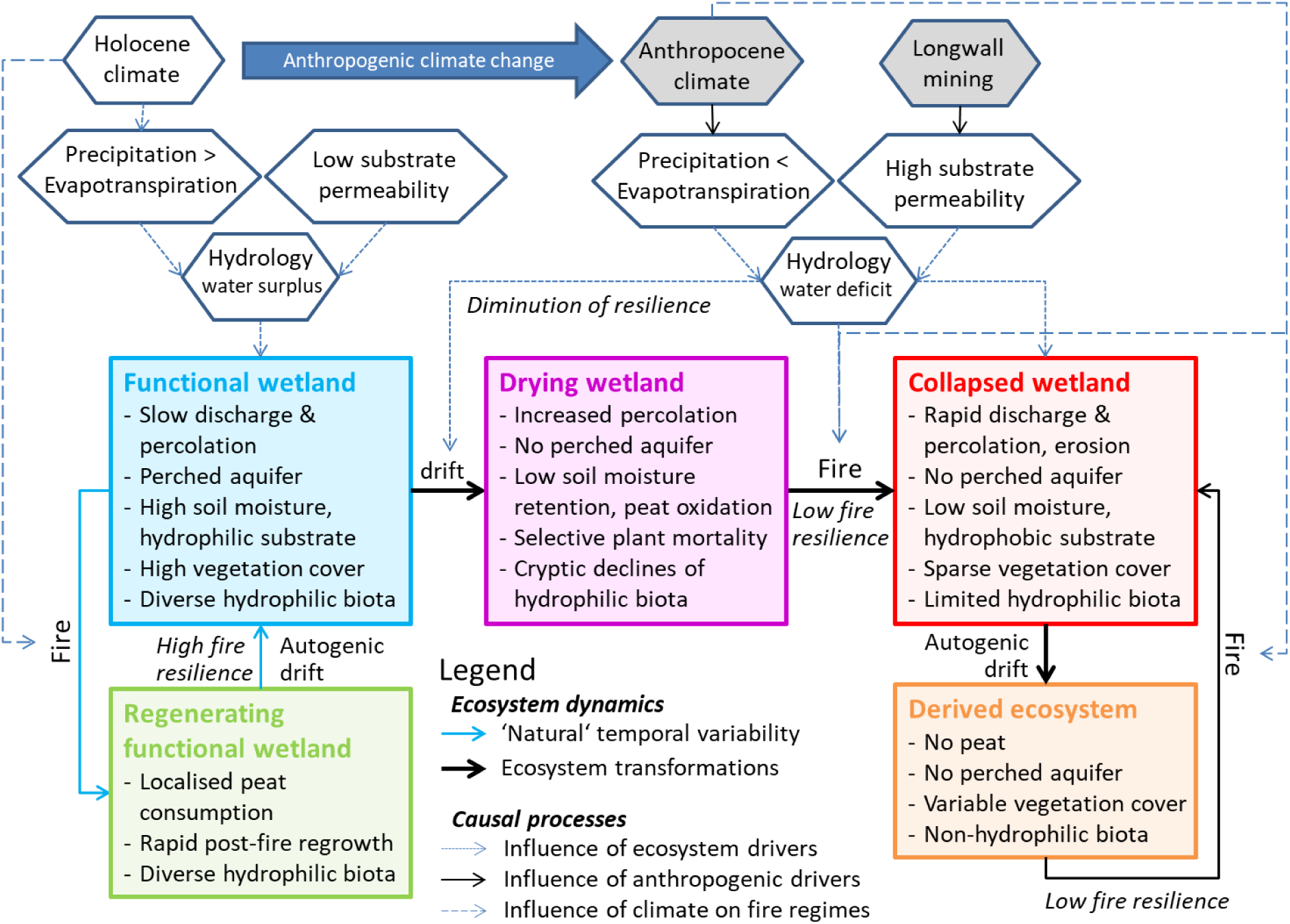
Conceptual model of ecosystem dynamics for groundwater-dependent, peat-accumulating wetland (upland swamps). Holocene hydrological conditions underpinned by climatic water surplus, impermeable substrates and flat terrain (not shown) facilitate development and maintenance of functional upland swamp wetlands (left) that are maintained by positive vegetation-soil-moisture feedbacks. Functional wetlands are resilient to recurring fires (perturbation regime) and return rapidly to mature state via autogenic drift when perturbed. Degradation of hydrological conditions by anthropogenic drivers, longwall mining (short time scales) or climate change (long time scales) causes transition of functional wetlands to a drying state (centre) and diminution of resilience to fire. Fire triggers a sudden transition of drying wetlands to a collapsed state (right), which drift autogenically to a derived ecosystem type that shares few of the features that characterise the functional wetlands.

We hypothesised that ecosystem collapse occurs when extreme drying of the peaty substrates diminishes resilience of the system to fire. The initial transition from a ‘Functional’ to a ‘Drying’ state (Fig. 2) may be cryptic, due to lagged persistence of dense vegetation as hydrological function declines, albeit with selective deaths of hydrophilic plant species. The persistence of dense vegetation, even when branches and shoots have partially senesced, maintains some moisture in the microclimate and limits entry of non-hydrophilic biota. We hypothesise that, due to reduced retention of substrate moisture (Mason, Krogh, Popovic, Glamore, & Keith, 2021), a Drying state is predisposed to severe consumption of peat and surface vegetation when ignited. This results in transition to a Collapsed state with further loss of hydrological function, mortality of standing hydrophilic plants and seedbanks, and consequent failure of the regenerative process that is typical of resilient and functional upland swamps (Fig. 2).

Longwall mining is the primary anthropogenic driver of wetland drying (Fig. 2). It involves complete extraction of a deep underground coal seam, allowing overburden rock to fall into the void, with associated upward shattering and cracking of bedrock, as well as warping, subsidence, upsidence and cracking at the surface (Booth, 2006) (Krogh, 2007). This alters hydrology by increasing the permeability of the substrate beneath the peatlands and by altering surface flows. We hypothesise that climate change is a secondary anthropogenic driver of wetland drying (Fig. 2), altering hydrology over longer time scales by increasing evapotranspiration relative to precipitation ((Keith, Rodoreda, & Bedward, Decadal change in wetland-woodland boundaries during the late 20th century reflects climatic trends, 2010); (Keith, Elith, & Simpson, Predicting distribution changes of a mire ecosystem under future climates, 2014)), subject to large interannual variability and regional climate cycles, with complex biochemical processes governing the breakdown of peat (Davidson & Janssens, 2006) (Limpens, et al., 2008).

### Experimental design and data collection

To determine whether mining-induced changes in hydrology weakened ecosystem resilience to fire, we compared the post-fire responses of wetlands that had underground mining beneath them with those of unmined reference wetlands (i.e. the two transitions labelled ‘Fire’ in Fig. 2). We tracked changes in resilience by monitoring soil moisture content. We predicted that contrasting post-fire responses of ecosystem resilience and collapse would be detectable through differences between reference peatlands and those influenced by longwall mining, respectively, in: i) peat retention/consumption; ii) post-fire vegetation structure and biomass; iii) plant species richness and composition; and iv) plant population processes (survival and reproduction).

We contextualised changes in ecosystem resilience by monitoring soil moisture annually during summer in three swamps affected by mine subsidence and three unmined reference swamps (Table 1). Measurements were taken at least 24 hours after rainfall in January or February each year, except in the summer of 2019-20, when they were taken in December towards the end of an extended drought and just prior to the bushfire, and the summer of 2021-22, when they were taken at the beginning of March after an extended period of rainfall. During each annual monitoring event we measured volumetric soil moisture content with a MP406 Soil Moisture Sensor Instant Reading Kit with a 6 cm probe, (ICT International, Armidale), which uses a standing wave oscillator to generate an electrical field to detect dielectric properties related to the moisture content of a substrate. We measured soil moisture at ten replicate sites (circular 1m radius) on the valley floor and sides within each swamp based on the mean of three randomly placed insertions of the probe.

To examine ecosystem responses to fire, we sampled five swamps affected by mine subsidence and five reference swamps, including the six sampled for soil moisture (Table S1). Within each swamp, we established one site on the valley floor and one site on the valley side to encompass variation in vegetation and hydrology. All sites were within a 10 × 5 km area and altitudinal range of 990 – 1120 m above sea level, and hence climatically similar. All sites were burnt on 16 December 2019 during extensive east Australian bushfires. Four of the sites (two mined and two reference) were also burnt in October 2013, one reference site was burnt in January 2003, and the remaining five sites had been unburnt since prior to 1980 (Table S1).

At each site, 20 m transects parallel to the swamp axis were sampled during 4-6 March 2020, approximately 10 weeks after fire, except sites CC and EW, which were sampled on 24 August 2020, While site BUD was only sampled in a second survey (below). We measured three metrics of fire severity (Keeley, 2009): i) scorch height, the lowest unscorched pre-fire plant tissues remaining on or adjacent to the transect, representing flame height; ii) the proportion of pre-fire foliage consumed or scorched on the transect for the woody and non-woody components of the vegetation, respectively (correlated with fire severity,; and iii) mean diameter of the smallest remaining twig or branch (n= 10) along the transect (a direct index of fire severity, (Whight & Bradstock, 1999).

To quantify vegetation structure, we measured the height (upper and lower bounds and mode), and visually estimated projective cover of live shrubs and graminoids in each transect. We also collected samples of above-ground live biomass (post-fire regrowth) from four 0.5 × 0.5m quadrats spaced at 5 m intervals along a line located parallel to, and 5 m from, each transect in similar vegetation to that along the transect. Samples were stored in clean paper bags, dried in an oven at 60°C until mass was stable and weighed in the laboratory.

We assessed peat loss during the fire by first visually estimating the percentage of the surface affected by peat consumption within each 20 × 1 m transect, and then measuring the maximum vertical distance from the current soil surface to the pre-fire soil level inferred from morphology and markings of exposed and charred root stocks of woody plants. We calculated an index of peat consumption from the product of these two values.

We identified and counted vegetatively resprouting individuals (R), dead remains of woody plants (D, i.e. individuals with no regenerating tissues) and seedling recruits (S) of all vascular plant taxa within each contiguous 1 × 1 m quadrat along the 20 m transects. Reconnaissance after survey in March 2020 indicated that some plants had delayed post-fire resprouting or germination responses (e.g. physiologically dormant species). We therefore resurveyed all plots in November 2020 to ensure that the species composition of regenerating vegetation was fully sampled across the first post-fire year. One reference site (BUD) was only sampled in the November survey. For all taxa, we estimated survival rates (R/(R+D)) and the maximum density of seedling recruitment (S) across the March and November surveys.

To examine the response of ecosystem fauna, we monitored abundance of a locally endemic lizard, *Eulamprus leuraensis* (Blue Mountains water skink), an Endangered species with a narrow hydrophilic environmental niche restricted to upland swamps. Skinks were trapped annually from 2013 to 2022 at the same six sites as those monitored for soil moisture (Appendix S1). At each site, each year on a day with temperature 20-35°C and no rainfall, we set 9 unbaited funnel traps and one pitfall trap c. 10 m apart, except in 2022 when 10 funnel traps were deployed (Gorissen, Greenlees, & Shine, 2017).

### Data analysis

We fitted a mixed log-linear model to examine temporal trends in soil moisture in relation to mining treatment and rainfall during the three months prior to soil measurement (rainfall data for November-January from Bureau of Meteorology, Lithgow (Cooewull) station 63226, 33.48° S 150.13° E, 900 m elevation, c. 5 km west of the study area). The model included an interaction term between mining influence (factor with two levels) and log-transformed time (t+1) in years (continuous), a main effects term for rainfall (continuous) and a random factor for site.

We used linear models to test the effects of underground mining and landform on fire severity (twig diameter), peat consumption, vegetation structure (height and cover of woody and non-woody plant strata, respectively), soil chemistry, plant biomass, plant survival and recruitment, and species richness (woody and non-woody). The models had two fixed factors (mining and landform) and an interaction term, with transects as replicates (N=20). Each of the factors had two levels (Mining influence: yes/no; Landform: valley floor/valley side). Models for all variables except plant survival and reproduction were fitted with normal error distributions and identity link functions. The data were log-transformed to improve residuals where assumptions for mean-variance, homoscedasticity and normality did not hold.

We compared species composition between longwall-mined and unmined reference swamps and landform types using a 2-factor multi-species generalised linear model of abundances (combined counts of resprouts and post-fire recruits) in the R package ‘mvabund’ (Wang, Naumann, Wright, & Warton, 2012). To accommodate variation in the timing of species responses through the first post-fire year, we took the maximum estimated abundance (combined counts of resprouts and seedlings) of that recorded in the March and November surveys for each species in each transect. Of 134 plant taxa recorded in either survey, we included 43 species in the multivariate model based on their occurrence in at least four (20%) of the 20 transects. We fitted the models using a negative binomial distribution with log link and checked residuals to confirm satisfactory representation of mean-variance relationships in the data. We removed terms with weak effects to simplify the multi-species model and carried out univariate tests to identify species that had the strongest responses to the remaining factors. The P-values were adjusted to control the family-wise error rate across species, using a resampling-based implementation of Holm’s step-down multiple testing procedure (Wang, Naumann, Wright, & Warton, 2012).

To visualise compositional relationships among samples, we constructed a Global Non-metric Multidimensional Scaling (GNMDS) ordination in 2 dimensions based on pairwise Bray-Curtis dissimilarity values in the R package ‘vegan’ (Okansen, et al., 2020). An Epsilon threshold was set at 0.95, to convert higher Bray-Curtis values to geodesic distances. Optimum (lowest stress) configurations were selected from 100 runs derived from random initial configurations with a maximum of 200 iterations for convergence to achieve convergence ratio 0.99999 from successive stress values. The two ordinations with the lowest stress values were compared with a Procrustes test and were found to be identical (r = 1, p = 0.001, permutations = 999).

For woody plant species with regenerative organs, we estimated plant survival as the proportion of individuals of species that had resprouted new foliage by the November 2020 survey. Transects (n=20) were replicates for each species. The proportion of survivors was analysed using linear mixed models with the same factorial design as above, but with species as added as a random independent variable and a binomial error distribution with logit link function to accommodate the bounded proportional values. Density of post-fire seedling recruits was analysed with same model, using a 1-factor multi-species generalised linear model of abundances to test effects of mining treatment.

We used a linear model with a Poisson error to compare trends in skink abundance by testing an interaction between mining treatment and time with thew summer rainfall tally as a covariable (see soil moisture model). All linear models were constructed in the R package, ‘lme4’ (R Core Team, 2020).

## Results

### Ecosystem drivers

Soil moisture levels declined in mined swamps, while they were maintained in unmined reference swamps (mining:time interaction t=13.42, P<<0.00001, Fig. 3). Declines in soil moisture began to occur soon after coal extraction and all three mined swamps fell below 50% soil moisture by 2018 (Fig 3). After the 2019 fire, soil moisture rarely exceeded 30% in mined swamps, whereas soil moisture remained in the range of 70-95% within unmined reference swamps throughout 2014-15 to 2021-22. Although prior rainfall had a positive effect on soil moisture (t=5.55, P<0.00001), and the trend varied among individual swamps (variance component 9.25, 95% CI: 4.52-18.95), differences between mined and unmined swamps were maintained through dry and wet years (92 mm in summer 2019-20, 560 mm in summer 2021-22) and appear to be irreversible.

**Figure 3.**
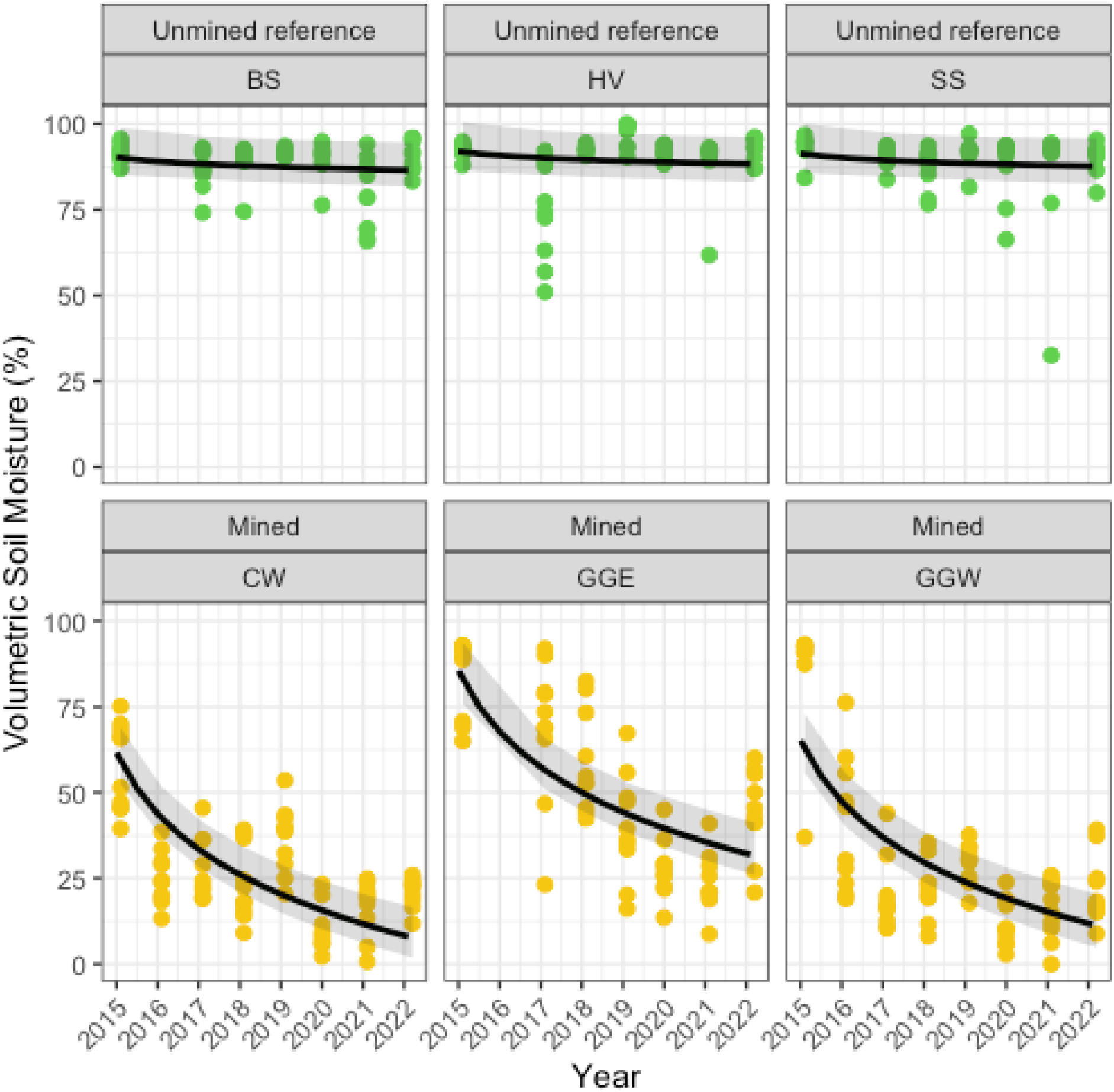
Trends in soil moisture indicative of ecosystem resilience in three unmined reference swamps (top) and three swamps exposed to underground longwall extraction of coal during 2014-17. Fitted lines are log-linear models Soil Moisture ∼Mining Treatment * log (time) + Rainfall. 95% confidence intervals were estimated using a nonparametric case bootstrap coupled with individual residual bootstrap for the range of rain values across all years.

Scorch height and % foliage consumed or scorched were uninformative indicators of fire severity, as all foliage in all 20 transects was completely consumed and scorch height always reached the highest branches of the tallest plants. Twig diameter data also showed no evidence of differences in fire severity among mining treatments (interaction & main effects t<1.72, P>0.2; Fig. 4a).

**Figure 4.**
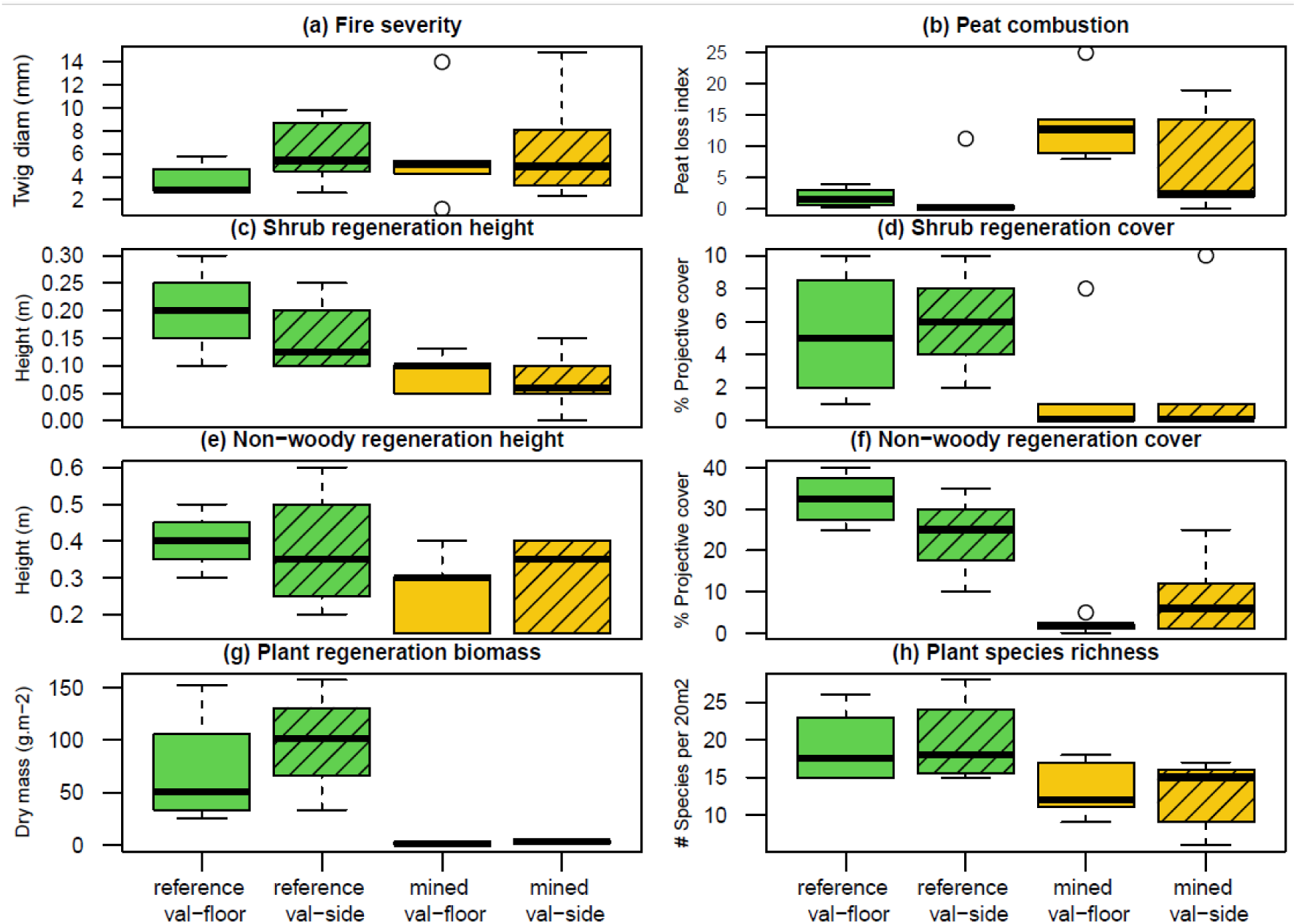
Comparative exposure of reference and mined swamps on their valley floors and sides to a) fire severity based on the minimum diameter of remaining shrub twigs or branches; and ecosystem responses based on b) peat combustion index; c) height of regenerating shrubs; d) cover of regenerating shrubs; e) height of regenerating non-woody plants; f) cover of regenerating non-woody plants; g) regenerating plant biomass; and h) plant species richness. All variables measured 10 weeks after the fire, except plant species richness, which was based on pooled species recorded 10 weeks (early autumn) and 11 months (late spring) after fire.

### Ecosystem state variables

Mining treatment had a very strong effect on peat loss (t=3.96, P=0.0019; Fig 4b), with the peat consumption index more than 5-fold greater in mined swamps compared to unmined reference swamps. Main effects landform types and their interaction with mining had no effect on peat loss (P=0.87 and P = 0.34, respectively).

The cover of shrubs 10 weeks post-fire in mined swamps was reduced by >95% of their cover in unmined reference swamps (t=5.81, P<0.0001;Fig. 4c), while cover of the non-woody ground layer was reduced by 20% (t=5.21, P=0.0002; Fig 4f), a difference that was sustained through the first post-fire year (Appendix S2). Longwall mining also reduced the height of regenerating shrubs by 50% (t=3.23, P=0.0072; Fig. 4d). Although shrub cover and height increased over time, the difference in height increased during the first post-fire year and the difference in cover was maintained (Appendix S2). The there was initially weak evidence of reduced height of regenerating non-woody vegetation in mined swamps (t=2.00, P=0.063; Fig. 4e), but within a year after fire it had grown taller in reference swamps (t=3.18, P=0.0051; Appendix S2). There was no effect of landform on any of the measured features of vegetation structure (Figs. 4c-f).

Biomass production of regenerating vegetation of mined swamps was 98% less than that in unmined reference swamps during the first 10 weeks after fire (t=7.06, P=0.00001, Fig. 4g). Substantial regrowth occurred over the ensuing months, but differences were maintained with biomass still 89% less in mined swamps than reference swamps 11 months after fire (Appendix S2). Biomass of post-fire regrowth was unaffected by landform (P>0.5).

Longwall mining initially reduced plant species richness of post-fire regrowth by 33% relative to unmined reference swamps (t=2.92, P=0.013, Fig. 4h), a pattern that was maintained 11 months after the fire, despite increases in richness across all treatments due to delayed germination of some species (Appendix S2). Richness was unaffected by landform irrespective of time since fire (main effects and interaction P>0.7 after 10 weeks, P>0.15 after 11 months).

A strong effect of longwall mining on plant species composition within the regenerating swamps was evident 10 weeks after fire, with samples from unmined reference sites clustering together in the ordination (stress=0.091), and mined sites dispersed more widely and strongly separated from the reference sites (Fig. 5). The segregation was maintained 11 months after fire (Appendix S2). The greater dispersion of mined samples is likely due to their lower richness and abundance of species. Landform had a secondary influence on species composition, with valley floor sites having more positive scores than corresponding valley side samples on first ordination axis (Fig. 5).

**Figure 5.**
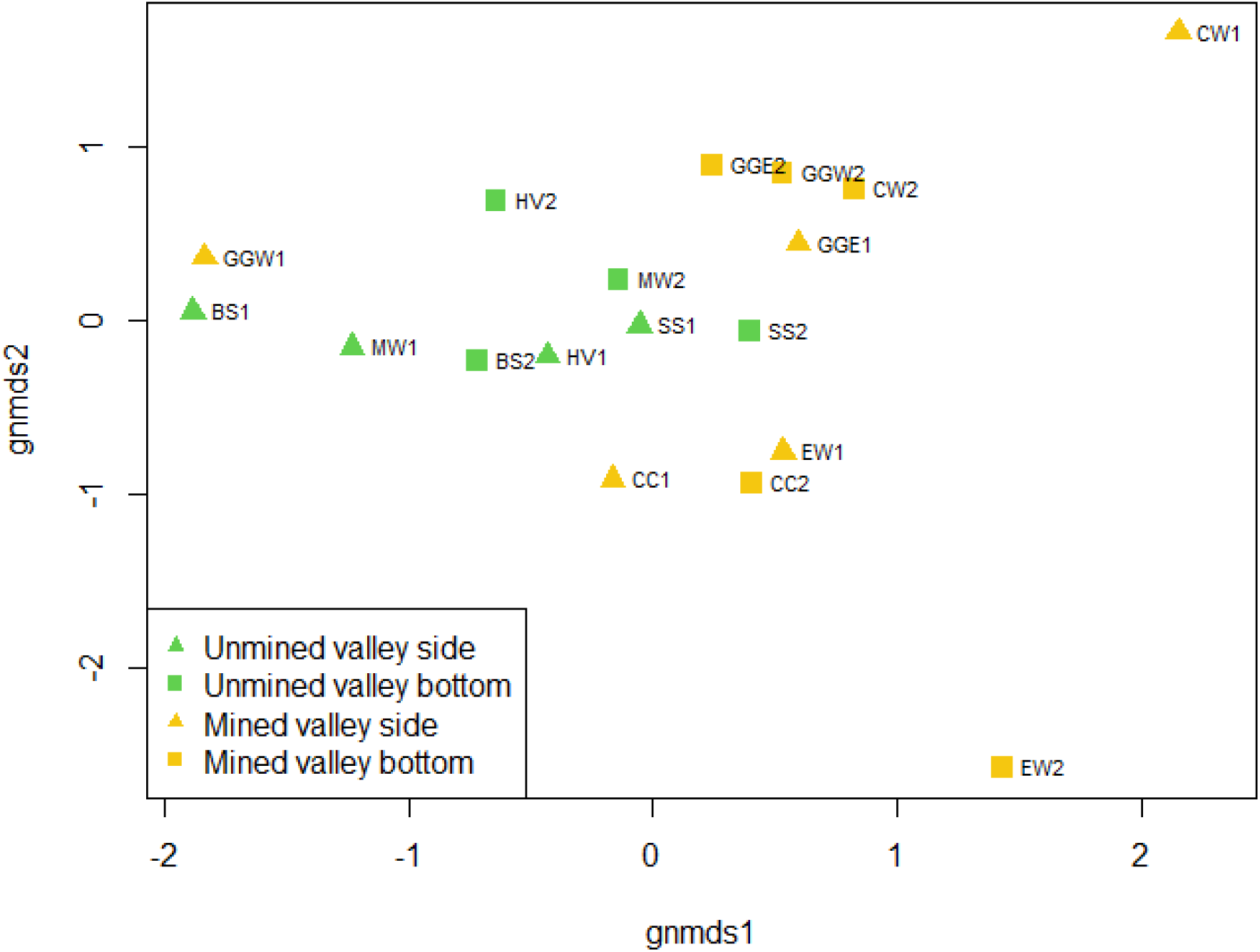
A global non-metric multidimensional scaling ordination showing differences in species composition between mined swamps and unmined reference swamps, as well as distinctions between landforms.

Overall, there was moderate evidence of differences in species composition between mined and unmined reference swamps (Wald=13.3, P=0.016). Of 43 plant species with sufficient occurrence for analysis (Appendix S2), five species were recorded exclusively in unmined reference swamps and one (*Eucalyptus oreades*) was recorded only in mined swamps (Appendix S2). There was strong evidence of differences in abundance between mining treatments in three species (P<<0.01), moderate evidence of differences in seven species (P<0.05), weak evidence of differences in a further four species (P<0.1), and weak evidence of differences due to mining on one of the landforms (interaction term P<0.1). Thus, just over half of the species examined showed some evidence of mining effects. Nine of the 43 species examined showed strong, moderate or weak evidence of differences in abundance between landform types (Appendix S2).

There was strong evidence of an interactive effect of mining treatment and landform on fire-related plant mortality (z=8.66, P<0.0001). Almost all (95%, with 95% confidence interval 86-99%) detectable pre-fire established plants were killed by fire on valley floors of mined swamps compared to only 2(1-7)% in unmined reference swamps, while fire mortality on valley sides of mined swamps was 66(42-84)% compared with 21(9-41)% in unmined reference swamps (Fig. 6). Mortality responses varied among the six species included in the model (variance 1.421, SD 1.192).

**Figure 6.**
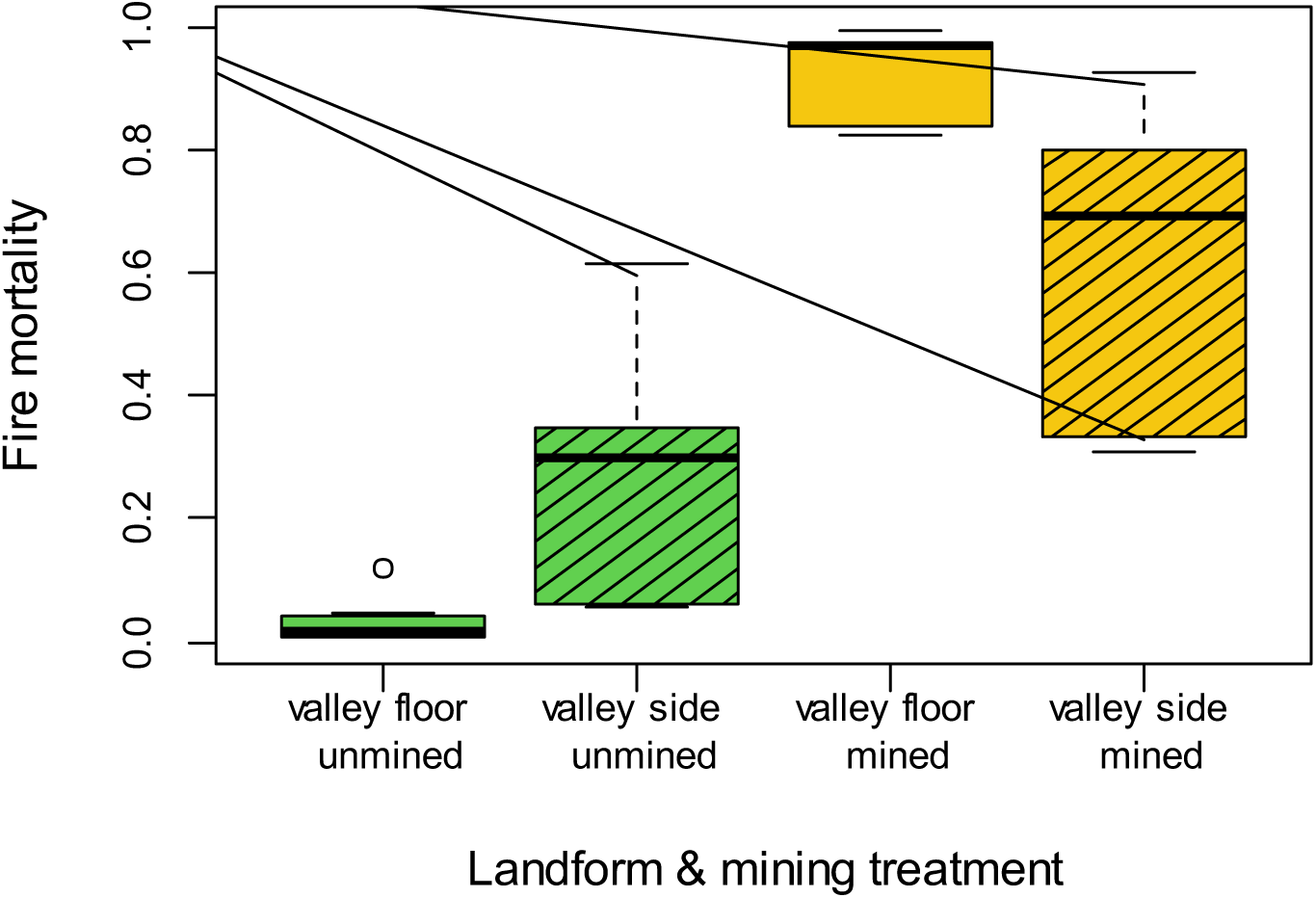
Variation in mortality rates of six detectable abundant resprouter plant species in relation to landform and mining treatments. Mortality estimates are those are predicted from a mixed binomial linear model with species as a random factor (see Methods text).

There was strong evidence that mining reduced post-fire seedling recruitment compared to unmined swamps (Deviance=74.9, P=0.004), but neither landform main effects (P=0.45) nor the interaction term (P=0.21) were important in the model of recruitment. Overall, the density of emerged post-fire seedlings in mined swamps was 11.9±1.4 m^-2^ compared with 56.0±6.9 m^-2^ in unmined reference swamps, with two species recorded only in reference swamps, one recorded only in mined swamps (*Eucalyptus oreades*), two species exhibiting moderate evidence of differences (P<0.05) and two species exhibiting weak evidence of differences (P<0.1) (Appendix S2).

Skink abundance declined approximately to zero in mined swamps but remained extant without strong trends in unmined reference swamps (mining:time interaction t=5.17, P<0.0001; Fig 7), with rainfall accounting for small component of interannual variation (t=2.57, P=0.010). No skinks were detected in any of the mined swamps in 2022.

**Figure 7.**
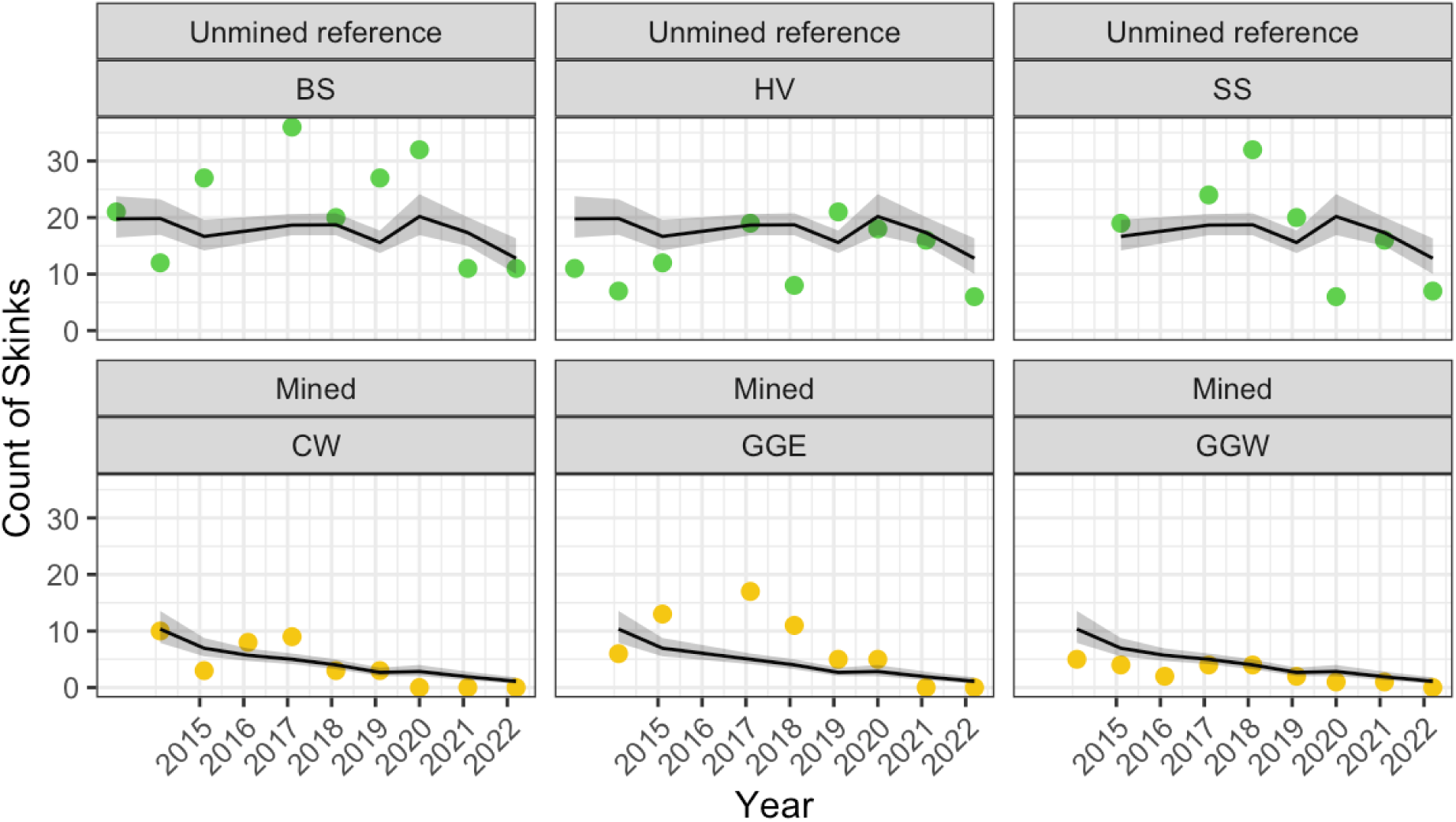
Trends in captures of *Eulamprus leuraensis* (Blue Mountains water skink), an obligate hydrophilic animal species, in three unmined reference swamps (top row) and three mined swamps (bottom row). 95% confidence intervals were estimated using a nonparametric case bootstrap coupled with individual residual bootstrap for the range of rain values across all years.

## Discussion

### Diminution of resilience precedes ecosystem collapse

Ecosystem responses to wildland fire differed markedly between swamps that had been exposed to underground mining and reference swamps that had not been exposed to mining. These differences in response were expressed in a wide range of ecosystem indicators, even though mined and unmined swamps experienced similar climatic conditions and similar fire severity (as estimated by several metrics). Relative to unmined reference swamps, mined swamps showed greater loss of peat via substrate combustion, reduced cover of regenerating vegetation (both woody and non-woody components), reduced height of regenerating shrubs, reduced biomass of regenerating vegetation, reduced post-fire plant species richness and abundance, altered plant species composition, increased mortality rates of woody plants, reduced post-fire seedling recruitment, and local extinction of a hydrophilic fauna species. (Turetsky, Donahue, & Benscoter, 2011) also recorded greater peat consumption in burnt fens that were drained by surface ditching, relative to burnt undrained reference fens. These differences indicate transformational changes in structure, composition and function consistent with ecosystem collapse in swamps exposed to underground mining (Fig. 8). The autogenic post-fire recovery process evident in unmined reference swamps was disrupted in the mined swamps, resulting in a new system with more slowly growing, sparser and shorter vegetation, with much reduced abundance of hydrophilic species that characterise functional swamps and entry of non-hydrophilic species including trees and non-native taxa.

**Figure 8.**
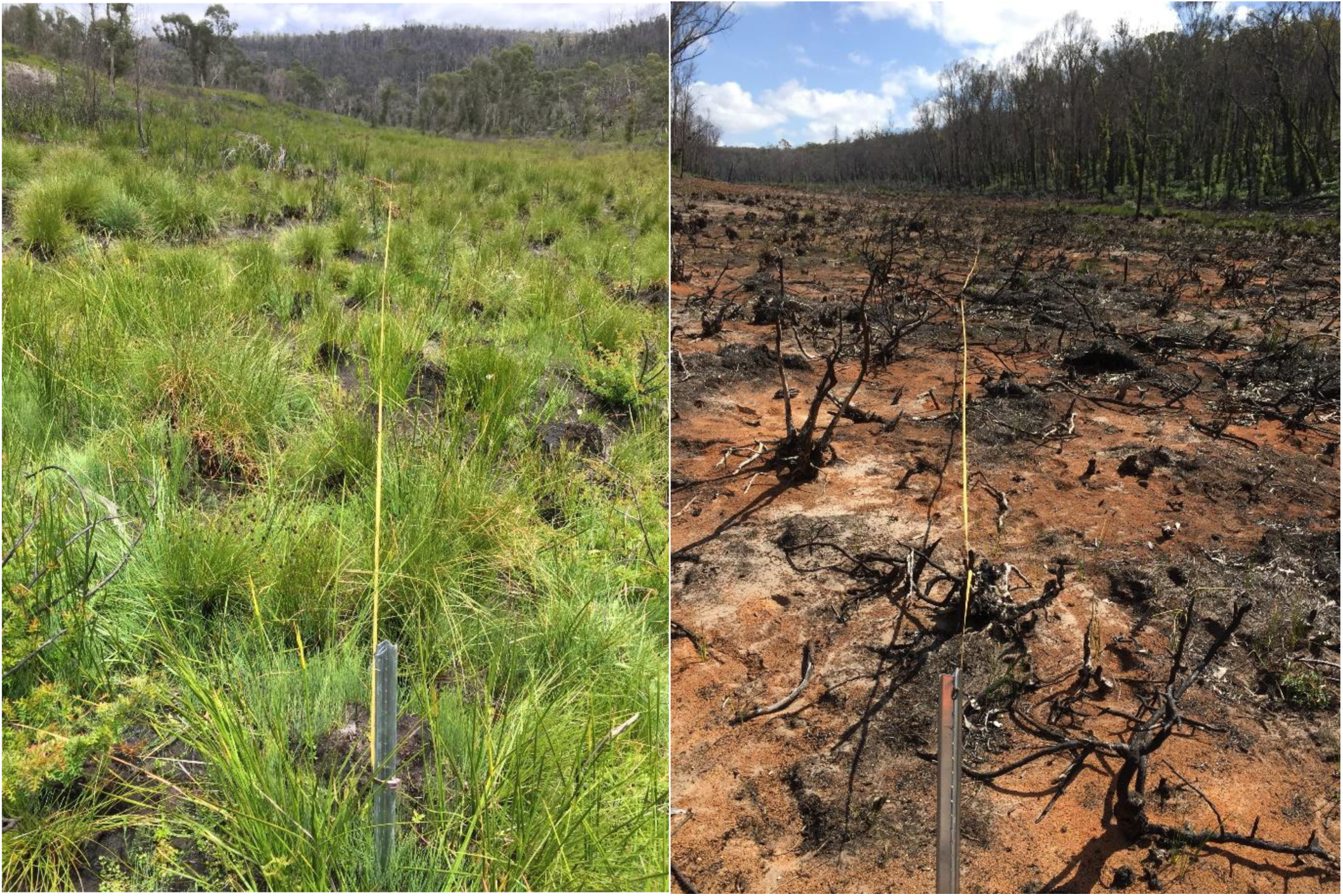
Resilient response to fire at Happy Valley (HV) reference swamp (left) and ecosystem collapse after underground mining and fire at Carne West swamp (right), both in the November 2020 survey, 11 months after bushfires in December 2019 [photo: David Keith, 25 November 2020].

The soil moisture data indicate that mined swamps underwent a change in hydrological regime soon after coal seam extraction but prior to the passage of fire. This prior change in a key ecosystem driver (groundwater hydrology) apparently predisposed the swamps to major impacts across the range of ecosystem indicators examined. We therefore conclude that our results support our model of ecosystem collapse (Fig. 1) in which a stressor (mining-related hydrological change) diminishes the resilience of an ecosystem to a perturbation (wildland fire). The post-fire responses of reference swamps suggest that, in the absence of the stressor, the ecosystems maintain resilience to the perturbation, enabling them to return to their ecological basin of attraction (Folke, et al., 2004).

Although we did not explicitly examine climate change in this study, our results have important implications for understanding its role in the sustainability of upland swamp ecosystems. Climate change may also be expected to diminish the resilience of the ecosystem to fires. In contrast to longwall mining, acts as an anthropogenic stressor on hydrology by increasing evapotranspiration relative to precipitation, rather than reducing substrate permeability.

Ecosystem responses to climate change may therefore be expected to be more delayed and more fluctuating (due to interannual weather cycles) than responses to longwall mining, but the transformation outcomes are likely to be similar, given the climatically marginal conditions for peat-forming ecosystems on mainland Australia. While climate change is a pervasive stressor subject to ecological lags and inertia of earth systems, longwall mining is a localised stressor, responsive to relatively short-term deterministic management decisions about where and how to extract coal. The two stressors act together (within mining footprints) or climate change acts alone (beyond mining footprints).Therefore, it should be possible to reduce and delay the total loss of resilience by managing the short-term stressor (longwall mining), allowing time for climate change mitigation measures to take effect on the long-term stressor (climatic water deficit).

### The consequences of ecosystem collapse

Concerns about the effect of underground longwall mining on groundwater and surface hydrology have been discussed for some time ((NSW-Scientific-Committee, 2005); (Krogh, 2007)). Knowledge was initially circumstantial, built on post-hoc observations of multiple independent drying and erosion events related to longwall mining (and associated subsidence, bedrock cracking and surface warping) that occurred in months and years prior to those events (Krogh, 2007). A review of remedial treatments found that none were successful in restoring the hydrology of impacted swamps (Commonwealth-of-Australia, 2014). Long term monitoring also showed no evidence of autogenic recovery of hydrological function over 5 years after mining (Mason, Krogh, Popovic, Glamore, & Keith, 2021).

The consequences of wetland ecosystem collapse are substantial for both biodiversity and ecosystem services. The Newnes Plateau Shrub Swamps belong to a global ecosystem functional group (TF1.6 Boreal, temperate and montane peat bogs; (Keith, Ferrer-Paris, Nicholson, & Kingsford, 2020) that has a very limited distribution in the southern hemisphere. On mainland Australia, these ecosystems are extremely restricted due to climatically marginal conditions for peat development. The Newnes Plateau Shrub Swamps belong to one of four such ecosystem types currently listed as Endangered Ecological Communities under national biodiversity legislation. The trends documented here indicate accelerating decline in its status. Biodiversity declines are also expressed at the species level, as the Newnes swamps host a range of locally endemic taxa and many that occupy narrow ecological niches that are predisposed to elevated extinction risks from hydrological change. Several plant and animal species essentially restricted to these ecosystems are already listed as threatened under national legislation (Krogh et al. in review). At landscape scales, peatland ecosystems are critical to supply of multiple ecosystem services, including carbon sequestration (Cowley & Fryirs, 2020), regulation of stream flow, mitigation of droughts and floods ((Young, 2017); (Cowley, Fryirs, & Hose, 2018)); supply of drinking water (Krogh, 2007); and maintenance of riverine and estuarine recreational waters. These values, together with the irreversibility of ecosystem collapse, underscore the need for prediction and early warning of ecosystem collapse (Scheffer, et al., 2009), as well as precautionary decision making to ensure ecologically sustainable development (Mason, Krogh, Popovic, Glamore, & Keith, 2021).

### Early diagnosis and risk reduction for collapse-prone ecosystems

(Dakos, Carpenter, van Nes, & Scheffer, 2015) identify conceptual, operational and methodological constraints on the ability of critical slowing down (CSD) indicators to signal early warning of ecosystem collapse. Conceptually, CSD indicators are most relevant to ecosystems exposed to slowly changing drivers that may generate abrupt changes in ecosystem state (Dakos, Carpenter, van Nes, & Scheffer, 2015). The collapse of Newnes swamps, however, involves strong stepwise change in stressor that sets in motion a rapid change in a hydrological driver (similar to mechanism (f) of (Dakos, Carpenter, van Nes, & Scheffer, 2015)), which is unlikely to be detected by CSD indicators. Even if CSD was detected from a hydrological time series, no management action could reverse hydrological decline after mining had initiated it (Commonwealth-of-Australia, 2014). Additionally, the methods used in our study to monitor a range of critical state variables are based on manual field sampling, which imposes large operational constraints on the acquisition of time series suitable for early detection CSD, even though efficient technologies exist to acquire dense time series for the hydrological driver. High density time series are practicable in some intensively used ecosystems such as fisheries or ecosystems in which remote sensing can provide an informative and sensitive state variable relevant to the mechanism of collapse, but resourcing is insufficient to monitor most of the world’s ecosystems on the ground at this level. In some ecosystem types, slow response rates of state variables may lengthen time series required to detect CSD signals, placing further operational constraints on their application (Bolt, B, van Nes, & Scheffer, 2021).

Early warning of collapse in many ecosystems therefore hinges on a mechanistic understanding of the processes that drive it. Even for CSD indicators, with theoretical foundations that confer generic applicability irrespective of precise mechanisms, a basic understanding of mechanisms is needed to assess whether they are conceptually appropriate and operationally and methodologically practicable. We suggest a four-point strategy for sustaining ecosystems: 1) diagnose potential mechanisms and likely causes of collapse; 2) reduce risks through actions that address causal drivers and maintain resilience; 3) slow and ameliorate expression of symptoms of collapse; and 4) restore ecosystem resilience.

Diagnosing the potential mechanisms of collapse has two components. The first is a qualitative diagnosis or ‘imaginative synthesis’ (Keith, Martin, McDonald-Madden, & Walters, 2011) to identify alternative ecosystem states, key components, and the causes, effects and dependencies of change. Experiential knowledge and qualitative diagrammatic models are powerful tools for such diagnostic synthesis (Keith, et al., 2015). This should reveal suitably proximal and sensitive state variables to examine ecosystem responses to the hypothesised stressors.

The second step of diagnosis is to quantify the thresholds in drivers and state variables that mark a state change. These threshold values are essential to inform timely preventative action. They may be estimated by probing the system experimentally, accumulating and synthesising case histories, or by drawing on indirect evidence from similar ecosystems as operational constraints permit ((Walter & Holling, 1990); (Keith, Martin, McDonald-Madden, & Walters, 2011)). For upland swamps, case histories show that collapse has occurred locally across a range of longwall mine designs that vary in panel width (Young, 2017), and that no-collapse outcomes may require mining exclusion zones (Mason, Krogh, Popovic, Glamore, & Keith, 2021).

As (Scheffer, et al., 2009)note, ecosystems in which collapse occurs in local units (such as in individual peatlands) or through reversible processes (such as in small lakes) offer more scope than singular systems to estimate thresholds by observation. For singular systems, such as ocean upwelling zones, extrapolation from models and learnings from related ecosystems will be the primary basis for threshold estimation (Bland, et al., 2018).

Risk reduction and impact avoidance is a preventative measure that is the only viable means of achieving ecosystem sustainability where collapse may be irreversible, at least over time scales practicable for ecosystem management. Based on active adaptive management approaches, alternative strategies should be deployed, monitored and evaluated to ‘learn by doing’ about the most effective risk reduction option ((Walter & Holling, 1990); (Williams, 2011)). Alternative risk reduction strategies that warrant investigation for our study system may involve mining exclusion zones of varying sizes and configurations, or different partial extraction designs with substantial seam retention (Mason, Krogh, Popovic, Glamore, & Keith, 2021). More generally, to reduce risk in other ecosystems managers can explore alternative fire regimes (e.g. in savannas and other fire-affected ecosystems), varied levels and patterns of harvest (e.g. in fisheries and other trophically regulated systems), varied pollution regulations (e.g. in stream or lake catchments), etc.

It may be possible to devise management strategies that delay or reduce the expression of symptoms of ecosystem decline, such as loss of biodiversity and function. By prolonging the time that ecosystems support biodiversity and supply services, these impact minimisation strategies can complement, rather than substitute for risk reduction strategies. Forestalling the expression of collapse may provide opportunities for lagged remedial measures take effect, for interventions to secure translocated or *ex situ* populations of affected species, or to develop and implement new restoration technologies. For example, the most severe symptoms of ecosystem collapse in Newnes swamps occurred abruptly after fire, even though declines in hydrological function, carbon sequestration and biodiversity were set in motion soon after coal seam extraction. Fire exclusion may not have avoided collapse of the peatlands, but would have slowed rates of carbon emission from peat and slowed rates of mortality of standing plants, their root mat and seed banks, prolonging the availability of material for translocation, the stability of sediments and the potential for regeneration if new technologies for restoring hydrological conditions were developed in future.

Finally, ecosystem restoration could be an effective strategy where the mechanism of ecosystem collapse is reversible. (Dakos, Carpenter, van Nes, & Scheffer, 2015)identify several mechanisms of collapse that may be reversible, including b) slow-fast cyclic transitions with negative feedbacks, c) stochastic resonance and e) long transient upon extreme events. Reversal may occur autogenically or in response to an environmental trigger, and may be promoted or accelerated by restoration management. However restoration techniques are still in their infancy in many ecosystem types, and there are relatively few examples of fully successful ecosystem restoration outcomes (Gann, et al., 2019). None have been demonstrated for peat-accumulating wetlands affected by underground mining ((Commonwealth-of-Australia, 2014)). At Newnes, for example, remediation measures failed to restore one of our study sites (East Wolgan swamp, (Young, 2017)). Preventative risk reduction will be a superior strategy for sustaining ecosystem biodiversity and function even where moderate uncertainties exist about the reversibility of collapse or the effectiveness of restoration techniques.

## Conclusions

An anthropogenic stressor may diminish the resilience of an ecosystem to recurring perturbations that otherwise promote characteristic temporal and spatial variability in the system. In our case study, longwall mining (an anthropogenic stressor) disrupted a vital ecosystem driver (hydrological function), diminishing the resilience of the resilience to a regular perturbation (wildland fire). A diverse range of ecosystem state variables encompassing structure, composition and structure showed that fire subsequently caused collapse of mine-affected swamps, while unmined reference swamps rapidly returned to their basin of attraction. Management implications. Avoidance of ecosystem collapse hinges on early diagnosis of mechanisms, and preventative risk reduction. In some cases, it may also be possible to delay or ameliorate symptoms of collapse or to restore resilience, although opportunities for the latter are limited where collapse is irreversible, as is the case for upland swamps studied here.

Ecosystem collapse may be viewed in the context of a broader socio ecological lens (Cumming & Peterson, 2017) in which economic norms have driven markets for coal and development of cost-efficient longwall extraction methods. A transition is underway to a clean energy future, driven by social norms and emergence of cost-effective alternative energy technology, and reducing demand for coal production. Implementing low-impact mine designs with suitable exclusion zones during the transitional phase, trading off marginal increments in the cost of coal extraction, should achieve large benefits in ecosystem risk reduction to maintain biodiversity and ecosystem services supported by upland swamp ecosystems. The alternative pathway, maximising coal production and its economic outputs prior to inevitable industry collapse, will cause irreversible ecosystem collapse and permanent loss of associated biodiversity and ecosystem services.

## Supporting information

EcosystemResilienceCollapse_20220326.pdf

EcosystemResilienceCollapse_20220326.pdf

