## Supplementary material for "Interactions between anthropogenic stressors and recurring perturbations mediate ecosystem resilience or collapse": EcosystemResilienceCollapse_20220326.pdf

**Table S1**. Field sample sites. * subset of sites used for soil moisture and skink monitoring.

| Swamp | Site | Easting | Northing | Mining treatment | Landform | Elevation (m) | Fire interval (years) |
|---|---|---|---|---|---|---|---|
| Budgary | BUD1 | 246057 | 6307835 | no | side | 997 | 17 |
| Budgary | BUD2 | 246050 | 6307850 | no | floor | 996 | 17 |
| Broad* | BS1 | 241955 | 6302758 | no | side | 1056 | >40 |
| Broad* | BS2 | 241924 | 6302797 | no | floor | 1053 | >40 |
| Marangaroo West | MW1 | 238812 | 6299342 | no | side | 1093 | 6.2 |
| Marangaroo West | MW2 | 238784 | 6299327 | no | floor | 1101 | 6.2 |
| Happy Valley* | HV1 | 241691 | 6297037 | no | side | 1074 | 6.2 |
| Happy Valley* | HV2 | 241683 | 6297046 | no | floor | 1075 | 6.2 |
| Sunnyside* | SS1 | 237790 | 6304112 | no | side | 1118 | >40 |
| Sunnyside* | SS2 | 237790 | 6304123 | no | floor | 1118 | >40 |
| Gang Gang East* | GGE1 | 240219 | 6302325 | yes | side | 1074 | 6.2 |
| Gang Gang East* | GGE2 | 240201 | 6302334 | yes | floor | 1074 | 6.2 |
| Gang Gang West* | GGW1 | 240091 | 6303038 | yes | side | 1068 | 6.2 |
| Gang Gang West* | GGW2 | 240091 | 6303023 | yes | floor | 1066 | 6.2 |
| Carne West* | CW1 | 239339 | 6303190 | yes | side | 1074 | >40 |
| Carne West* | CW2 | 239327 | 6303196 | yes | floor | 1076 | >40 |
| Carne Central | CC1 | 241187 | 6302715 | yes | side | 1067 | >40 |
| Carne Central | CC2 | 241160 | 6302690 | yes | floor | 1069 | >40 |
| East Wolgan | EW1 | 235961 | 6304150 | yes | side | 1108 | >40 |
| East Wolgan | EW2 | 235945 | 6304140 | yes | floor | 1105 | >40 |
