## Supplementary material for "Interactions between anthropogenic stressors and recurring perturbations mediate ecosystem resilience or collapse": EcosystemResilienceCollapse_20220326.pdf

### Appendix S2. Statistical models fitted to ecosystem drivers and state variables.

#### Ecosystem drivers

##### Soil moisture

Linear mixed effects model

We modelled variation in Soil Moisture (volumetric %), in relation to three explanatory variables: log(Years since 2014); a factor with two levels for mined and unmined sites (using treatment contrasts), and rainfall in the three months over late spring - summer (November-January) just prior to sampling. We added one random effect of Site and a constant variance function with two different levels for mined and unmined sites. The variance is fixed to one for the control level (Unmined reference), and estimated for the treatment level (Mined). The model is:

```
mdl000 <- lme(SoilMoisture ~ logYears * Mine + Rain, random = ~ 1 | Site,
        weights = varIdent(form= ~ 1 | Mine), data=newnsoil)
```

Linear mixed-effects model fit by REML

| AIC <dbl> | BIC <dbl> | logLik <dbl> |
|---|---|---|
| 3402.231 | 3434.834 | -1693.116 |

Random effects:

Formula: ~1 | Site

```
        (Intercept) Residual
StdDev:    9.251101 8.303085
```

Variance function:

Structure: Different standard deviations per stratum

Formula: ~1 | Mine

Parameter estimates:

Unmined reference Mined

1.000000 1.765437

Fixed effects: SoilMoisture ~ logYears * Mine + Rain

| | Value <chr> | Std.Error <chr> | DF <chr> | t-value <chr> | p-value <chr> |
|---|---|---|---|---|---|
| (Intercept) | 86.39378 | 5.590549 | 431 | 15.453542 | 0.0000 |
| logYears | -1.77496 | 0.917966 | 431 | -1.933576 | 0.0538 |
| MineMined | -20.43813 | 8.034454 | 4 | -2.543811 | 0.0637 |
| Rain | 0.01962 | 0.003537 | 431 | 5.546988 | 0.0000 |
| logYears:MineMined | -23.87648 | 1.779146 | 431 | -13.420193 | 0.0000 |

Correlation:

```
                   (Intr) logYrs MinMnd Rain
logYears           -0.214
MineMined          -0.680  0.163
Rain               -0.146 -0.132 -0.004
logYears:MineMined  0.119 -0.508 -0.311  0.009
```

Standardized Within-Group Residuals:

```
       Min         Q1        Med         Q3        Max
-6.6708225 -0.5213518  0.1804146  0.5683773  2.4939164
```

Number of Observations: 440

Number of Groups: 6

Approximated 95% confidence intervals for the variance components:

Hide

intervals(mdl201,which='var-cov')

Approximate 95% confidence intervals

```
 Random Effects:
  Level: Site
                    lower     est.    upper
sd((Intercept)) 4.516312 9.251101 18.94973

 Variance function:
         lower     est.   upper
Mined 1.539334 1.765437 2.02475
attr(,"label")
[1] "Variance function:"

 Within-group standard error:
   lower     est.    upper
7.527907 8.303085 9.158087
```

### Fire severity

Fire severity was represented by diameter of the largest remaining twigs 1-2 m above ground (scorch height and % cover scorched/consumed were uninformative). We fitted a 2-factor model to examine the effect of mining treatment and landform (valley floor cf. valley side). We logged the twig diameter to improve homogeneity of varance.

Model:

```
Logtwig~Mine*Val
           Df Sum Sq Mean Sq F value Pr(>F)
Mine        1 0.2651  0.2651   0.963  0.349
```

```
Val           1 0.2340  0.2340    0.850  0.378
Mine:Val      1 0.4731  0.4731    1.719  0.219
Residuals    10 2.7518  0.2752
```

### Ecosystem state variables

#### Vegetation structure

We measured height and cover of woody and non-woody plant strata in March 2020 (10 weeks after fire) and November 2020 (11 months after fire), and fitted a 2-factor model to examine the effect of mining treatment and landform (valley floor cf. valley side).

##### *Shrub cover*

**March 2020 survey**

```
log(Shbcov+1)~Mine*Val

Residuals:
     Min       1Q   Median       3Q      Max 
-0.94605 -0.19928  0.01589  0.32363  0.75870 

Coefficients:
                Estimate Std. Error t value Pr(>|t|)    
(Intercept)      1.63919    0.26720   6.135 0.000111 ***
Mineyes         -1.60742    0.40816  -3.938 0.002783 ** 
Valside          0.20789    0.37788   0.550 0.594297    
Mineyes:Valside  0.05493    0.57722   0.095 0.926063    
```

Simplified model: LogShbcov ~ Mine

Residuals:

```
    Min      1Q  Median      3Q     Max
-1.04999 -0.16318 -0.06787  0.30292  0.65476
```

Coefficients:

```
          Estimate Std. Error t value Pr(>|t|)
(Intercept)  1.7431     0.1781  9.786 4.52e-07 ***
Mineyes     -1.5800     0.2721  -5.807 8.38e-05 ***
```

Random factor for swamps not important and omitted.

**November 2020 survey**

```
LogShbcovNov20 ~ Mine * Val

Residuals:
     Min       1Q   Median       3Q      Max 
-1.15475 -0.71670 -0.03985  0.54838  1.48372 

Coefficients:
                Estimate Std. Error t value Pr(>|t|)    
(Intercept)       1.8479     0.3616   5.110 0.000105 ***
Mineyes          -1.0290     0.5114  -2.012 0.061358 .  
Valside           0.3768     0.5114   0.737 0.471875    
Mineyes:Valside  -0.4790     0.7233  -0.662 0.517208    

Simplified model:  LogShbcovNov20 ~ Mine
```

```
Residuals:
     Min       1Q   Median       3Q      Max 
-1.34318 -0.76779 -0.08253  0.40333  1.53480 

Coefficients:
            Estimate Std. Error t value Pr(>|t|)    
(Intercept)   2.0363     0.2454   8.297 1.45e-07 ***
Mineyes      -1.2685     0.3471  -3.655  0.00181 ** 
---
```

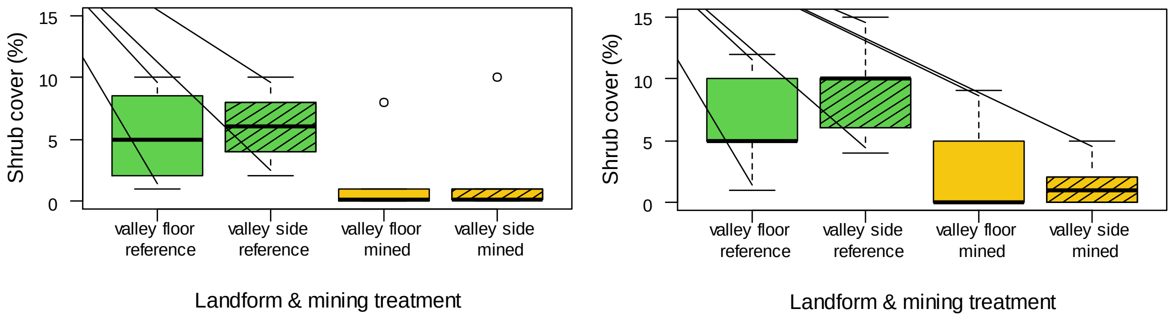


**Figure S2.1**. Shrub cover in March 2020 (left) and November 2020 (right)

*Shrub height*

**March 2020 survey**

```
LogShbhgt~Mine*Val

Residuals:
     Min       1Q   Median       3Q      Max 
-0.62123 -0.32078  0.07192  0.33689  0.58585 

Coefficients:
                Estimate Std. Error t value Pr(>|t|)    
(Intercept)     -1.68136    0.22006  -7.640 1.76e-05 ***
Mineyes         -0.76482    0.33615  -2.275   0.0462 *  
Valside         -0.29079    0.31121  -0.934   0.3721    
Mineyes:Valside  0.03306    0.47538   0.070   0.9459    

Simplified model: LogShbhgt~Mine

Residuals:
     Min       1Q   Median       3Q      Max 
-0.47583 -0.42069  0.07347  0.27246  0.62278 

Coefficients:
            Estimate Std. Error t value Pr(>|t|)    
(Intercept)  -1.8268     0.1516 -12.051 4.61e-08 ***
Mineyes      -0.7483     0.2315  -3.232   0.0072 ** 
```

**November2020 survey**

```
ShbhgtNov20 ~ Mine * Val

Residuals:
   Min     1Q Median     3Q    Max 
-0.130 -0.100 -0.015  0.100  0.220 

Coefficients:
                Estimate Std. Error t value Pr(>|t|)    
(Intercept)      0.30000    0.05303   5.657 3.57e-05 ***
```

```
Mineyes         -0.20000    0.07500  -2.667    0.0169 *
Valside         -0.02000    0.07500  -0.267    0.7931
Mineyes:Valside  0.01000    0.10607   0.094    0.9261
---

Simplified model: ShbhgtNov20 ~ Mine

Residuals:
    Min      1Q  Median      3Q     Max
-0.1400 -0.0950 -0.0175  0.1050  0.2100

Coefficients:
            Estimate Std. Error t value Pr(>|t|)
(Intercept)  0.29000    0.03545   8.180 1.78e-07 ***
Mineyes     -0.19500    0.05014  -3.889  0.00107 **
---
```

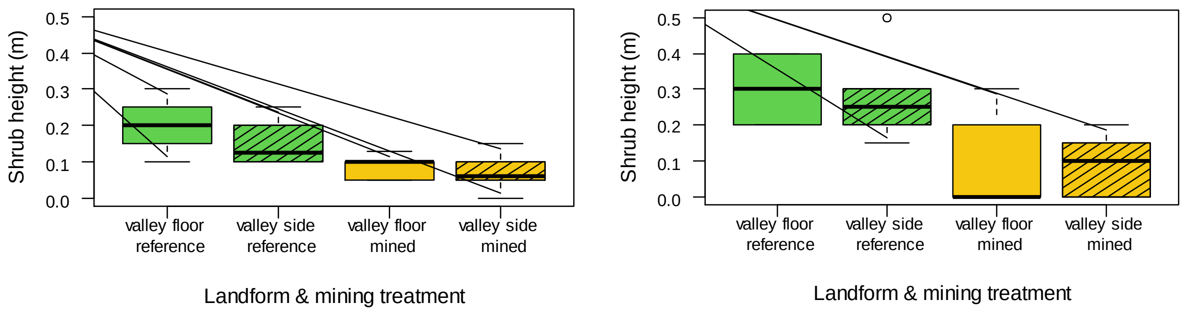


**Figure S2.2**. Shrub height in March 2020 (left) and November 2020 (right)

#### *Non-woody plant cover*

**March 2020 survey**

```
HrbcovMar20~Mine*Val

Residuals:
    Min      1Q  Median      3Q     Max
-13.750  -2.575   1.250   2.450  11.250

Coefficients:
                Estimate Std. Error t value Pr(>|t|)
(Intercept)       32.500      3.730   8.713 5.53e-06 ***
Mineyes          -29.800      5.698  -5.230 0.000384 ***
Valside           -8.750      5.275  -1.659 0.128166
Mineyes:Valside   14.717      8.058   1.826 0.097759 .

Simplified model: HrbcovMar20~Mine

Residuals:
    Min      1Q  Median      3Q     Max
-18.125  -3.125  -1.683   6.235  11.875

Coefficients:
            Estimate Std. Error t value Pr(>|t|)
(Intercept)   28.125      2.819   9.976 3.68e-07 ***
Mineyes      -22.442      4.307  -5.211 0.000218 ***
```

**November 2020 survey**

```
HrbcovNov20 ~ Mine * Val

Residuals:
   Min     1Q Median     3Q    Max
-14.00  -6.00  -1.91   6.75  16.00

Coefficients:
```

```
                  Estimate Std. Error t value Pr(>|t|)
(Intercept)         49.000      4.135  11.850 2.47e-09 ***
Mineyes            -44.180      5.848  -7.555 1.16e-06 ***
Valside            -13.000      5.848  -2.223   0.0410 *
Mineyes:Valside     19.180      8.270   2.319   0.0339 *
---
```

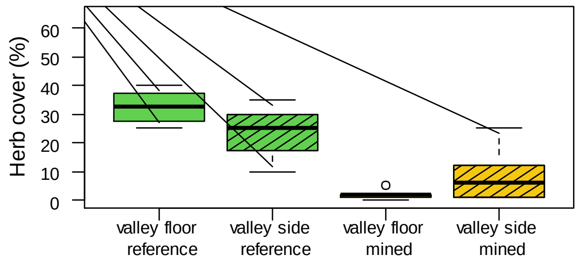


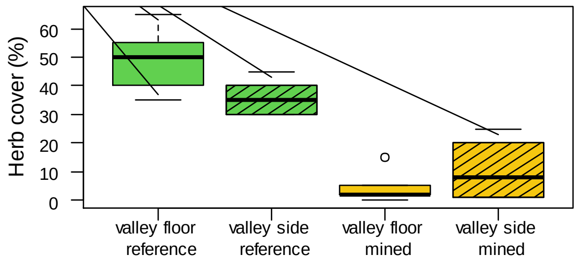


**Figure S2.3**. Non-woody plant cover in March 2020 (left) and November 2020 (right)

#### *Non-woody plant height*

**March 2020 survey**

```
HrbhgtMar20 ~ Mine * Val)

Residuals:
    Min      1Q  Median      3Q     Max 
-0.1750 -0.1075  0.0125  0.0900  0.2250 

Coefficients:
                Estimate Std. Error t value Pr(>|t|)
(Intercept)      0.40000    0.06289   6.360 1.77e-05 ***
Mineyes         -0.14000    0.08438  -1.659    0.119
Valside         -0.02500    0.08894  -0.281    0.783
Mineyes:Valside  0.05500    0.11933   0.461    0.652
---

Simplified model: HrbhgtMar20~Mine

Residuals:
    Min      1Q  Median      3Q     Max 
-0.1875 -0.1156  0.0125  0.1031  0.2125 

Coefficients:
            Estimate Std. Error t value Pr(>|t|)
(Intercept)  0.38750    0.04193   9.242 8.13e-08 ***
Mineyes     -0.11250    0.05625  -2.000   0.0628 .
---
```

**November 2020 census**

```
HrbhgtNov20~Mine*Val

Residuals:
    Min      1Q  Median      3Q     Max 
-0.1600 -0.0700  0.0000  0.0625  0.1900 

Coefficients:
                  Estimate Std. Error t value Pr(>|t|)
(Intercept)      4.000e-01  4.664e-02   8.577 2.22e-07 ***
```

```
Mineyes          -1.400e-01  6.595e-02  -2.123    0.0497 *
Valside           5.586e-17  6.595e-02   0.000    1.0000
Mineyes:Valside  -6.206e-17  9.327e-02   0.000    1.0000
---
```

```
Simplified model: HrbhgtNov20~Mine

Residuals:
    Min      1Q  Median      3Q     Max
-0.1600 -0.0700  0.0000  0.0625  0.1900

Coefficients:
            Estimate Std. Error t value Pr(>|t|)
(Intercept)  0.40000    0.03109  12.865 1.63e-10 ***
Mineyes     -0.14000    0.04397  -3.184  0.00514 **
```

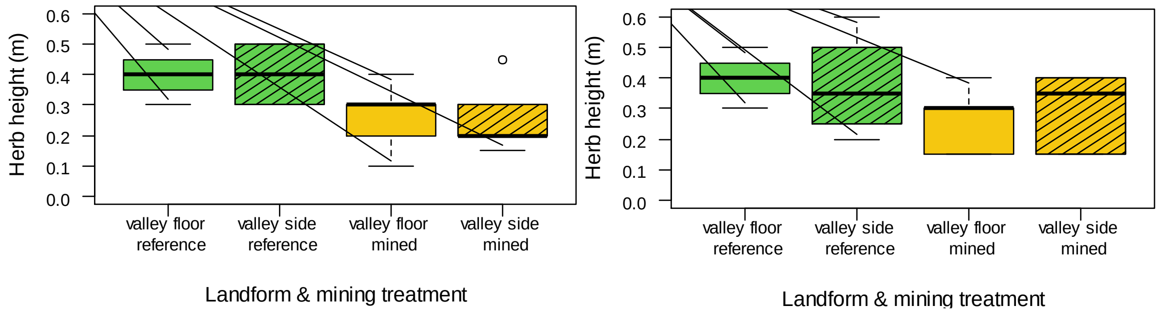


**Figure S2.4**. Height of non-woody plants in March 2020 (left) and November 2020 (right)

### Plant regeneration biomass

**March 2020 survey**

```
LogBmass~Mine*Val

Residuals:
    Min      1Q  Median      3Q     Max
-2.1063 -0.6614  0.1662  0.6010  1.2528

Coefficients:
                Estimate Std. Error t value Pr(>|t|)
(Intercept)       4.0156     0.5231   7.677 1.69e-05 ***
Mineyes          -4.4837     0.7990  -5.612 0.000224 ***
Valside           0.4341     0.7397   0.587 0.570295
Mineyes:Valside   0.8236     1.1300   0.729 0.482801
---

Simplified model: LogBmass~Mine

Residuals:
    Min      1Q  Median      3Q     Max
-2.7351 -0.4376  0.2904  0.7507  1.2049

Coefficients:
            Estimate Std. Error t value Pr(>|t|)
(Intercept)   4.2327     0.3777  11.207 1.03e-07 ***
Mineyes      -4.0719     0.5769  -7.058 1.32e-05 ***
---
```

**November 2020 survey**

```
LogBmassNov20~Mine*Val
```

```
Residuals:
    Min      1Q  Median      3Q     Max 
-3.3198 -0.2925  0.0877  0.5942  2.3412 

Coefficients:
                Estimate Std. Error t value Pr(>|t|)    
(Intercept)       5.8858     0.7606   7.738 8.52e-07 ***
Mineyes          -3.2170     1.0757  -2.991  0.00865 ** 
Valside          -0.2798     1.0757  -0.260  0.79806    
Mineyes:Valside  -0.3029     1.5212  -0.199  0.84470    
---

Simplified model: LogBmassNov20 ~ Mine

Residuals:
    Min      1Q  Median      3Q     Max 
-3.6112 -0.1819  0.0881  0.7720  2.4015 

Coefficients:
            Estimate Std. Error t value Pr(>|t|)    
(Intercept)   5.7458     0.5128  11.205 1.51e-09 ***
Mineyes      -3.3684     0.7252  -4.645 0.000201 ***
---
```

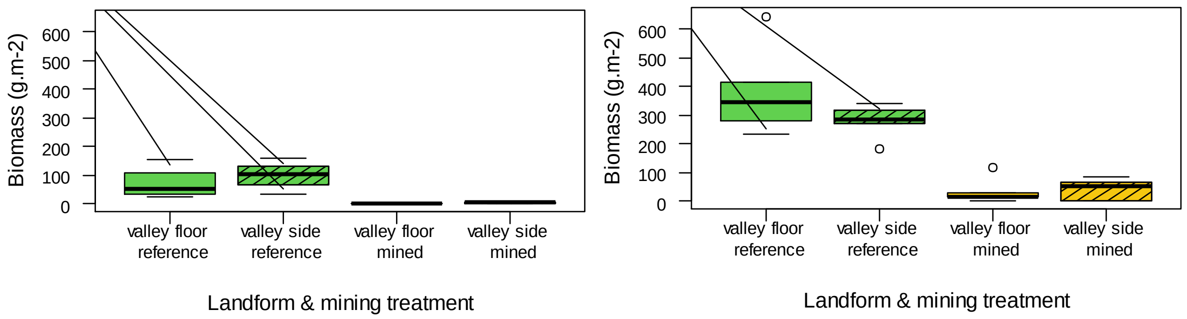


**Figure S3.5**. Plant regenerative biomass post-fire in March 2020 (left) and November 2020 (right)

### Species richness

**March 2020 survey**

```
Spprich20 ~ Mine * Val)

Residuals:
    Min      1Q  Median      3Q     Max 
-5.0000 -4.0000  0.1667  1.3333  9.0000 

Coefficients:
                Estimate Std. Error t value Pr(>|t|)    
(Intercept)       19.000      2.466   7.703 1.64e-05 ***
Mineyes           -8.333      3.768  -2.212   0.0514 .  
Valside            1.000      3.488   0.287   0.7802    
Mineyes:Valside    2.000      5.328   0.375   0.7152    
---
```

Simplified model

```
formula = Spprich20 ~ Mine)

Residuals:
    Min      1Q  Median      3Q     Max 
```

```
-4.5000 -3.4167 -0.6667  2.2500  9.5000

Coefficients:
            Estimate Std. Error t value Pr(>|t|)    
(Intercept)   19.500      1.642  11.876 5.43e-08 ***
Mineyes       -7.333      2.508  -2.924   0.0128 *  
---
```

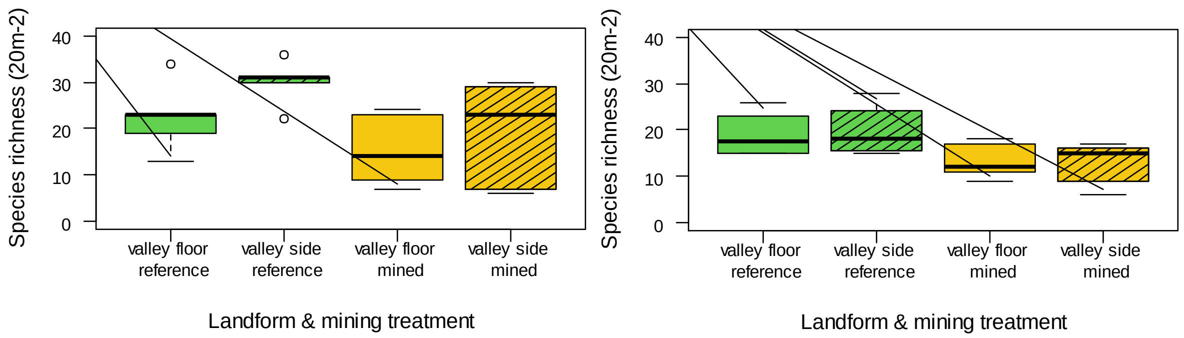


**Figure S2.6**. Plant species richness in March 2020 (left) and November 2020 (right)

### Plant species composition

We estimated the abundance (density of individuals) of all vascular plant taxa in 20m2 transects in March 2020 and November 2020 after fire in December 2019. 43 species occurred in at least 20 of the transects and were suitable for analysis.

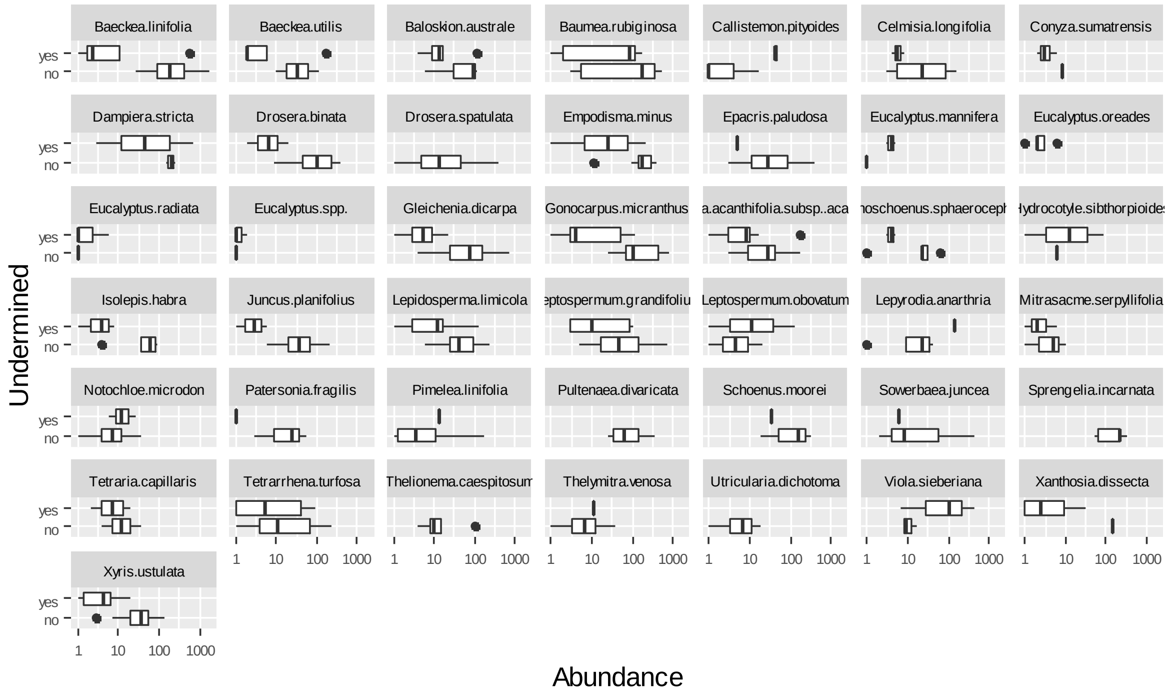


**Figure 2.7**. Plant species abundance (number of individuals per 20m$^2$) in swamps exposed to underground mining relative to unmined reference swamps.

A multivariate model was fitted to the 43 plant species with occurrence in at least four of 20 transects, with Wald tests for differences between mining treatments, landforms and their interaction.

```
>modfreq_gt3_counts3 <- manyglm(newnmv_gt3_counts ~ mine*val, family="negative_binomial")
>anova(modfreq_gt3_counts3, test = "wald", p.uni="unadjusted", cor.type = "shrink")
```

**Table S2.1**. Multivariate analysis of variance on plant species abundances (nomenclature follows PlantNet, https://plantnet.rbgsyd.nsw.gov.au/search/simple.htm).

| | Mine Wald | Mine P | | Val Wald | Val P | | Mine:Val Wald | Mine:Val P | |
|---|---|---|---|---|---|---|---|---|---|
| Multivariate (all 43 spp) | 13.37 | 0.016 | * | 11.44 | 0.122 | | 8.14 | 0.063 | + |
| *Baeckea linifolia* | 1.5 | 0.123 | | 3.309 | 0.041 | * | 1.748 | 0.171 | |
| *Baeckea utilis* | 0.195 | 0.756 | | 1.993 | 0.227 | | 0.425 | 0.268 | |
| *Baloskion australe* | 1.207 | 0.098 | + | 1.323 | 0.195 | | 1.025 | 0.257 | |
| *Baumea rubiginosa* | 1.3 | 0.118 | | 2.357 | 0.072 | + | 0.474 | 0.602 | |
| *Callistemon pityoides* | 0.903 | 0.326 | | 0.788 | 0.495 | | 0.905 | 0.112 | |
| *Celmisia longifolia* | 2.024 | 0.136 | | 0.037 | 0.954 | | 0.49 | 0.206 | |
| *Conyza sumatrensis* | 0.214 | 0.609 | | 1.502 | 0.09 | + | 0.032 | 0.362 | |
| *Dampiera stricta* | 0.206 | 0.56 | | 0.062 | 0.871 | | 0.001 | 0.794 | |
| *Drosera binata* | 2.953 | 0.016 | * | 1.859 | 0.132 | | 0.127 | 0.642 | |
| *Drosera spatulata* | 0.059 | 0.87 | e | 2.649 | 0.1 | + | 0.005 | 0.479 | |
| *Empodisma minus* | 2.446 | 0.004 | ** | 0.884 | 0.347 | | 0.842 | 0.403 | |
| *Epacris paludosa* | 4.243 | 0.001 | *** | 0.689 | 0.647 | | 0.034 | 0.369 | |
| *Eucalyptus mannifera* | 1.092 | 0.197 | | 0.431 | 0.568 | | 0.032 | 0.36 | |
| *Eucalyptus oreades* | 0.048 | 0.831 | e | 0.307 | 0.741 | | 0 | 0.736 | |
| *Eucalyptus radiata* | 1.513 | 0.131 | | 1.357 | 0.118 | | 0.045 | 0.123 | |
| *Eucalyptus* spp. | 1.216 | 0.085 | + | 0.452 | 0.48 | | 0.03 | 0.291 | |
| *Gleichenia dicarpa* | 4.034 | 0.001 | *** | 0.486 | 0.688 | | 1.727 | 0.14 | |
| *Gonocarpus micranthus* | 2.731 | 0.019 | * | 0.267 | 0.807 | | 0.48 | 0.699 | |
| *Grevillea acanthifolia* subsp. *acanthifolia* | 1.044 | 0.475 | | 0.927 | 0.572 | | 2.631 | 0.1 | + |
| *Gymnoschoenus sphaerocephalus* | 2.399 | 0.051 | + | 0.902 | 0.42 | | 0.035 | 0.608 | |
| *Hydrocotyle sibthorpioides* | 1.537 | 0.179 | | 0.058 | 0.869 | | 0.004 | 0.521 | |
| *Isolepis habra* | 2.659 | 0.039 | * | 0.686 | 0.497 | | 0.026 | 0.946 | |
| *Juncus planifolius* | 20523 | 0.086 | + | 2.004 | 0.121 | | 1.336 | 0.106 | |
| *Lepidosperma limicola* | 1.364 | 0.132 | | 3.309 | 0.015 | * | 2.447 | 0.057 | + |
| *Leptospermum grandifolium* | 2.053 | 0.115 | | 1.012 | 0.49 | | 1.375 | 0.3 | |
| *Leptospermum obovatum* | 0.878 | 0.451 | | 2.557 | 0.115 | | 0.034 | 0.275 | |
| *Lepyrodia anarthria* | 0.234 | 0.683 | | 1.304 | 0.334 | | 0.044 | 0.396 | |
| *Mitrasacme serpyllifolia* | 0.506 | 0.564 | | 0.664 | 0.539 | | 0.043 | 0.496 | |
| *Notochloe microdon* | 0.351 | 0.638 | | 0.149 | 0.834 | | 1.243 | 0.109 | |
| *Patersonia fragilis* | 2.481 | 0.045 | * | 0.065 | 0.918 | | 0.007 | 0.611 | |
| *Pimelea linifolia* | 1.98 | 0.206 | | 3.351 | 0.031 | * | 0.026 | 0.277 | |
| *Pultenaea divaricata* | 0.059 | 0.735 | e | 0.526 | 0.334 | | 0.002 | 0.672 | |

| | Mine Wald | Mine P | | Val Wald | Val P | | Mine:Val Wald | Mine:Val P | |
|---|---|---|---|---|---|---|---|---|---|
| *Schoenus moorei* | 10229 | 0.107 | | 0.062 | 0.917 | | 0.004 | 0.524 | |
| *Sowerbaea juncea* | 2.074 | 0.172 | | 0.072 | 0.882 | | 0.007 | 0.663 | |
| *Sprengelia incarnata* | 0.061 | 0.811 | e | 0.15 | 0.797 | | 0 | 0.812 | |
| *Tetraria capillaris* | 0.036 | 0.975 | | 0.445 | 0.694 | | 1.162 | 0.075 | + |
| *Tetrarrhena turfosa* | 1.377 | 0.211 | | 3.262 | 0.041 | * | 1.807 | 0.121 | |
| *Thelionema caespitosum* | 0.055 | 0.805 | e | 1.031 | 0.34 | | 0.002 | 0.534 | |
| *Thelymitra venosa* | 2.3 | 0.037 | * | 3.329 | 0.016 | * | 0.03 | 0.506 | |
| *Utricularia dichotoma* | 0.051 | 0.79 | e | 0.585 | 0.265 | | 0.001 | 0.721 | |
| *Viola sieberiana* | 1.655 | 0.235 | | 0.114 | 0.867 | | 0.738 | 0.134 | |
| *Xanthosia dissecta* | 0.677 | 0.564 | | 2.357 | 0.087 | + | 0.037 | 0.334 | |
| *Xyris ustulata* | 3.3 | 0.011 | * | 0.537 | 0.604 | | 1.766 | 0.139 | |

### Plant mortality caused by fire

Fire-killed dead remains of six woody species were reliably detectable and identifiable, and sufficiently abundant to fit a binomial model of mortality to examine differences between mining treatments and landforms.

```
Generalized linear mixed model fit by maximum likelihood
  (Laplace Approximation) [glmerMod]
 Family: binomial  ( logit )
Formula: cbind(Kill, Respr) ~ Mine * Val + (1 | Sppf)
   Data: data

     AIC      BIC   logLik deviance df.resid
   327.0    337.7   -158.5    317.0       57

Scaled residuals:
    Min      1Q  Median      3Q     Max
-6.8624 -0.9155 -0.0207  0.7420  8.2104

Random effects:
 Groups Name        Variance Std.Dev.
 Sppf   (Intercept) 1.421    1.192
Number of obs: 62, groups:  Sppf, 6

Fixed effects:
                Estimate Std. Error z value Pr(>|z|)
(Intercept)      -3.7973     0.6072  -6.254 4.00e-10 ***
Mineyes           6.8124     0.5237  13.009  < 2e-16 ***
Valside           2.4465     0.3807   6.426 1.31e-10 ***
Mineyes:Valside  -4.7887     0.5528  -8.662  < 2e-16 ***
---
Signif. codes:  0 '***' 0.001 '**' 0.01 '*' 0.05 '.' 0.1 ' ' 1

Correlation of Fixed Effects:
            (Intr) Mineys Valsid
Mineyes     -0.419
Valside     -0.567  0.654
Mineys:Vlsd  0.393 -0.931 -0.671
```

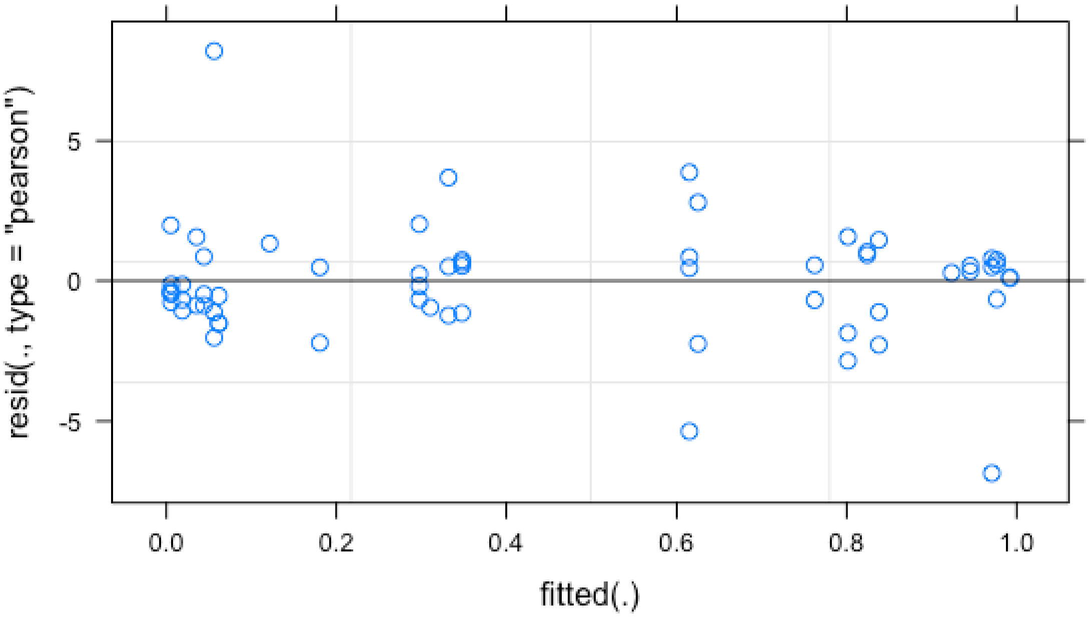


```
# Predicted probabilities of cbind(Kill, Respr)

# Val = floor

Mine | Predicted |       95% CI
--------------------------------
no   |      0.02 | [0.01, 0.07]
yes  |      0.95 | [0.86, 0.99]

# Val = side

Mine | Predicted |       95% CI
--------------------------------
no   |      0.21 | [0.09, 0.41]
yes  |      0.66 | [0.42, 0.84]

Adjusted for:
* Sppf = 0 (population-level)
```

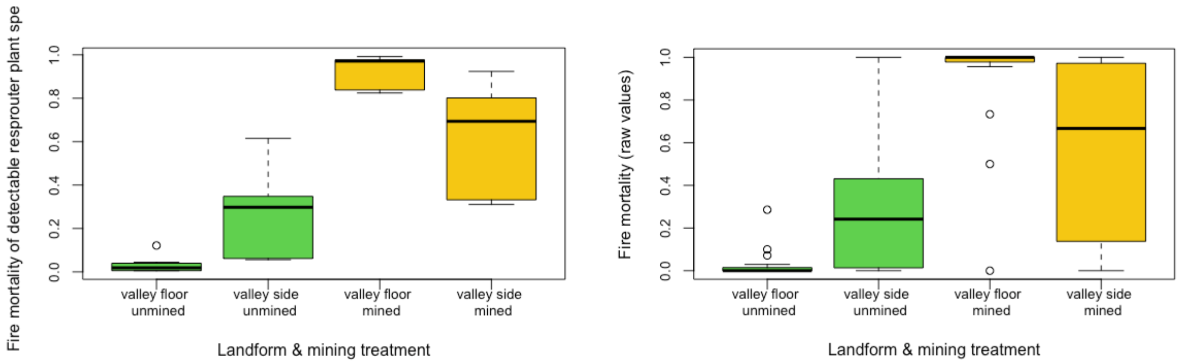


**Figure S2.8**. Plant mortality caused by fire – modelled estimates (left) and observed values (right)

### Post-fire plant recruitment

Multivariate model for 24 plant species with occurrence in at least four of 20 transects with Wald test for differences between mining treatments.

```
>modfreq_gt3_counts3 <- manyglm(newnmv_gt3_counts ~ mine,
family="negative_binomial")
>anova(newnmv_recruit_gt3_counts3, test = "wald", p.uni="unadjusted", cor.type =
"shrink")
```

**Table S2.2**. Abundance of post-fire seedling recruits in relation to mining treatment

| Species | Mine Wald | Mine P | |
|---|---|---|---|
| Multivariate (all 24 spp) | 6.7 | 0.093 | + |
| *Baeckea linifolia* | 2.821 | 0.03 | * |
| *Baumea rubiginosa* | 0.172 | 0.747 | |
| *Celmisia longifolia* | 0.200 | 0.745 | |
| *Conyza sumatrensis* | 0.214 | 0.611 | |
| *Dampiera stricta* | 0.047 | 0.872 | |
| *Drosera spatulata* | 0.050 | 0.864 | e |
| *Epacris paludosa* | 0.060 | 0.863 | |
| *Eucalyptus oreades* | 0.048 | 0.831 | e |
| *Eucalyptus radiata* | 1.513 | 0.112 | |
| *Eucalyptus* spp. | 1.216 | 0.105 | + |
| *Gonocarpus micranthus* | 0.021 | 0.021 | * |
| *Grevillea acanthifolia* subsp. *acanthifolia* | 0.785 | 0.591 | |
| *Gymnoschoenus sphaerocephalus* | 2.399 | 0.051 | + |
| *Hydrocotyle sibthorpioides* | 1.537 | 0.179 | |
| *Isolepis habra* | 2.229 | 0.081 | + |
| *Juncus planifolius* | 1.524 | 0.192 | |
| *Lepidosperma limicola* | 1.339 | 0.123 | |
| *Leptospermum grandifolium* | 1.537 | 0.225 | |
| *Mitrasacme serpyllifolia* | 0.397 | 0.612 | |
| *Notochloe microdon* | 0.905 | 0.270 | |
| *Pimelea linifolia* | 1.734 | 0.206 | |
| *Leptospermum obovatum* | 0.878 | 0.451 | |
| *Pultenaea divaricata* | 0.048 | 0.733 | e |
| *Sprengelia incarnata* | 0.061 | 0.794 | e |
| *Tetrarrhena turfosa* | 10229 | 0.107 | |
| *Viloa sieberiana* | 1.615 | 0.175 | |
| *Xanthosia dissecta* | 0.91 | 0.503 | |

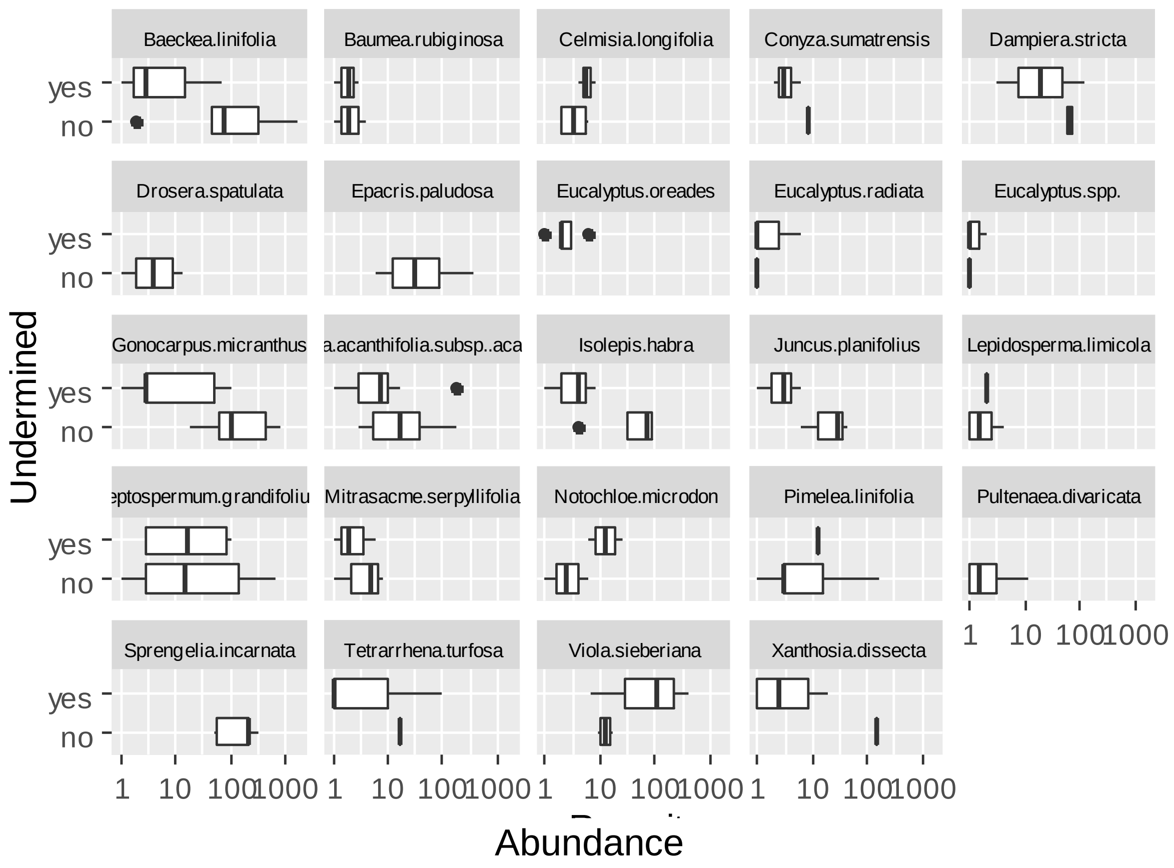


**Figure S2.9**. Maximum density of post-fire seedling recruits recorded across March and November 2020 surveys.

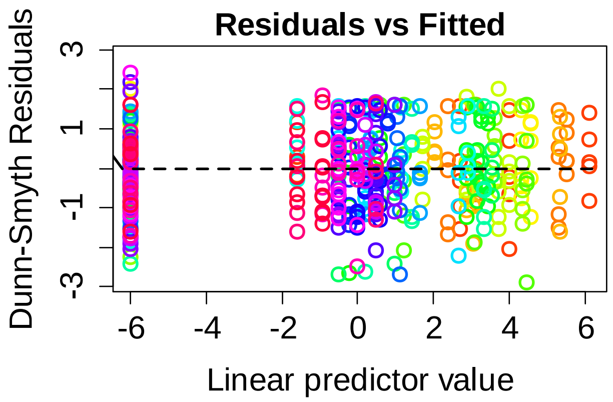


### Abundance of hydrophilic skink species

*Eulamprus leurensis* (Blue Mountains water skink) is found only in upland swamp habitats within a small geographic range in the upper Blue Mountains, west of Sydney.

Counts of captured skinks were modelled against the interaction of mining treatment and time with rainfall in the 3 months prior to survey as a covariable.

```
mdl <- glm(SkinkCount ~ Years * Mine + Rain , skink,family=poisson)
Deviance Residuals:
    Min       1Q   Median       3Q      Max
-3.7164  -1.6627  -0.4636   1.1129   4.1961
```

```
Coefficients:
                  Estimate Std. Error z value Pr(>|z|)    
(Intercept)      3.1238005  0.1028599  30.369  < 2e-16 ***
Years           -0.0052478  0.0182331  -0.288 0.773487    
MineMined       -0.6286620  0.1618002  -3.885 0.000102 ***
Rain            -0.0009499  0.0003703  -2.565 0.010314 *  
Years:MineMined -0.2219797  0.0429670  -5.166 2.39e-07 ***
---

    Null deviance: 432.39  on 50  degrees of freedom
Residual deviance: 168.92  on 46  degrees of freedom
AIC: 361.18
```

Residuals

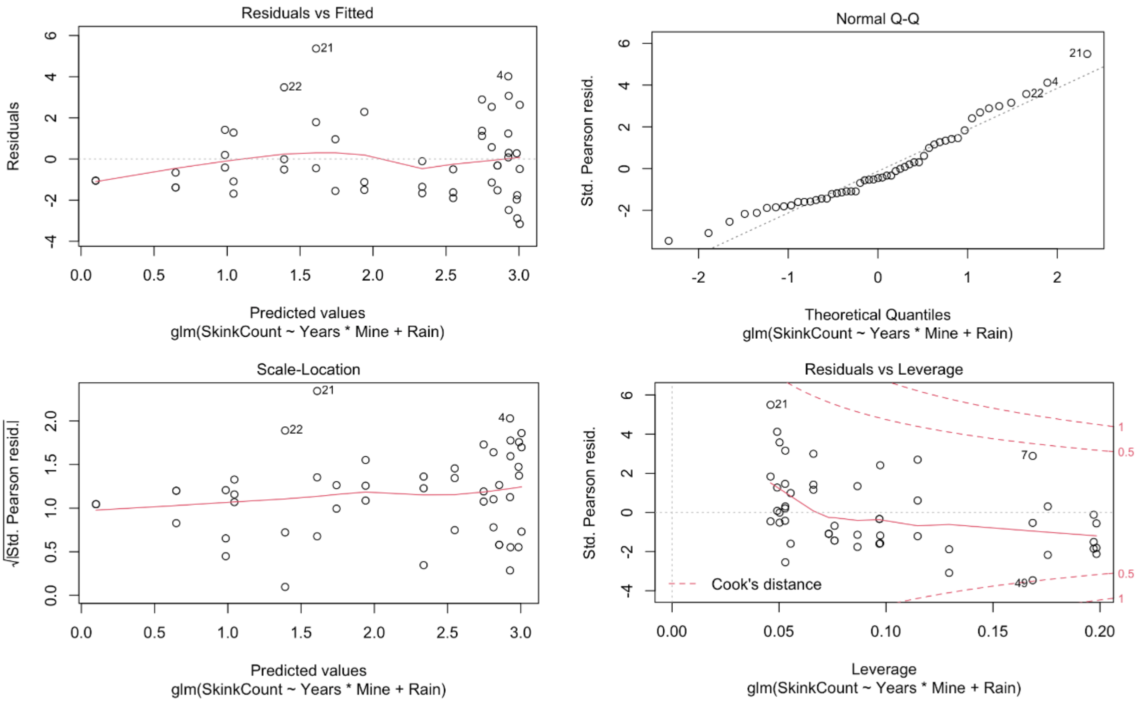

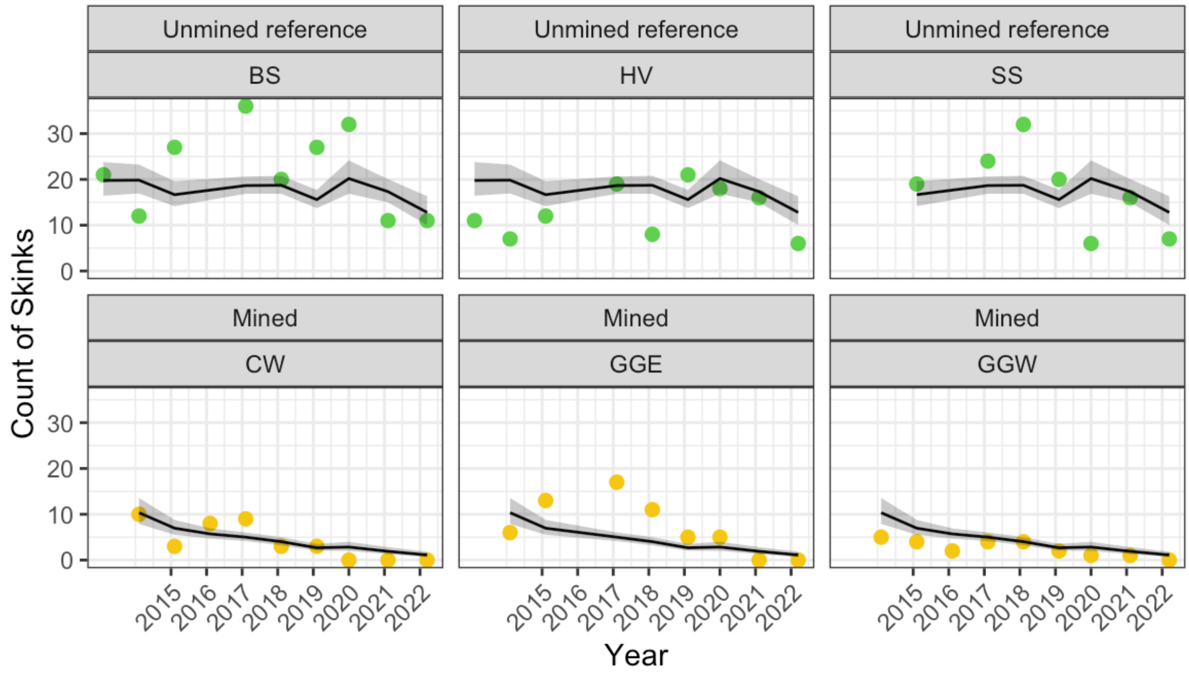


**Figure S.10**. Predicted skink counts and 95% confidence intervals with the measured values of rain.

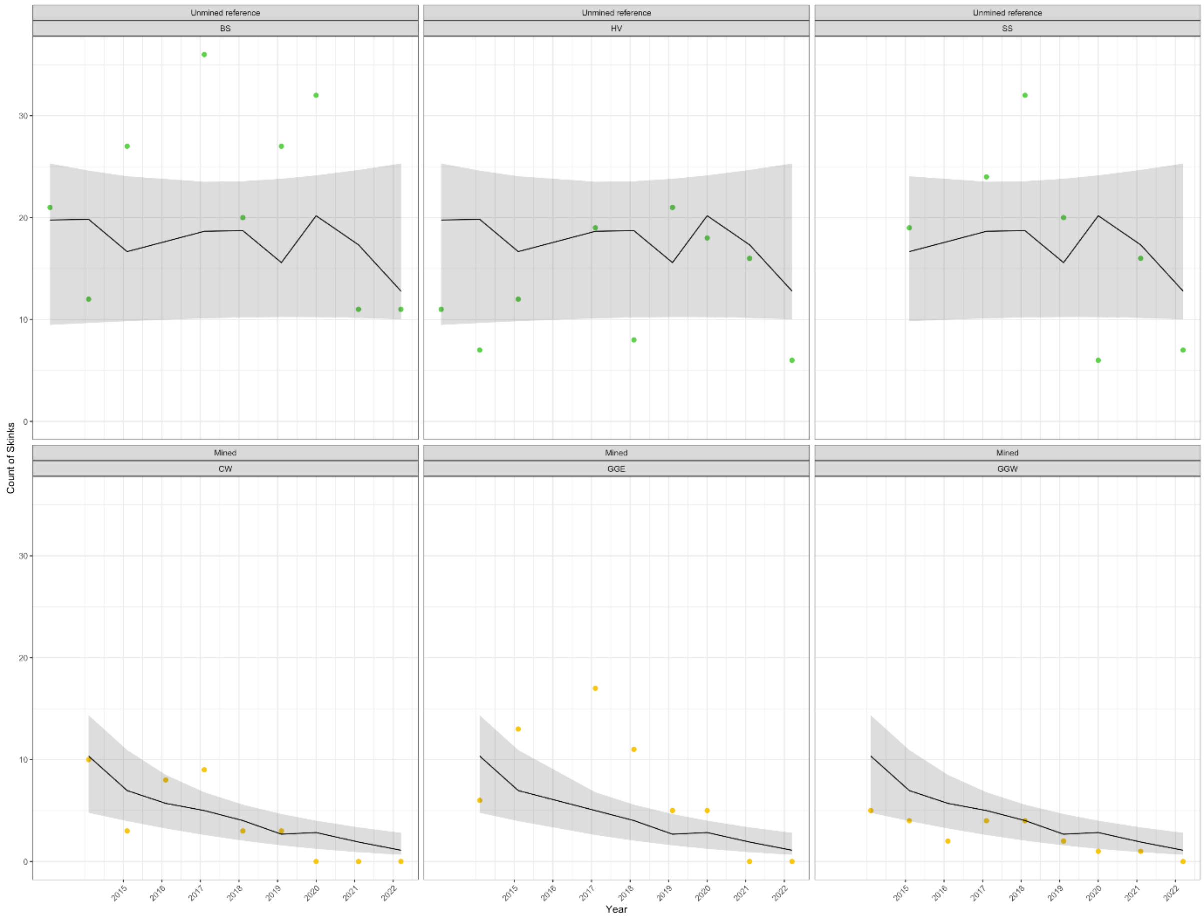


**Figure S.11**. Predicted skink counts and 95% confidence intervals with the range of values of rain
